# MHC class Ia molecules facilitate MCK2-dependent MCMV infection of macrophages and virus dissemination to the salivary gland

**DOI:** 10.1101/2022.10.05.510922

**Authors:** Berislav Bošnjak, Elisa Henze, Yvonne Lueder, Kim Thi Hoang Do, Alaleh Rezalofti, Christiane Ritter, Anja Schimrock, Stefanie Willenzon, Hristo Georgiev, Lea Fritz, Melanie Galla, Karen Wagner, Martin Messerle, Reinhold Förster

**Author notes:** Corresponding authors: Berislav Bošnjak, Institute of Immunology, Hannover Medical School, Carl-Neuberg Straße 1, 30625 Hannover, Germany, Reinhold Förster, Institute of Immunology, Hannover Medical School, Carl-Neuberg Straße 1, 30625 Hannover, Germany. This authors equally contributed (Berislav Bošnjak, Elisa Henze).

## Abstract

Murine cytomegalovirus (MCMV) infection of macrophages relies on MCMV-encoded chemokine 2 (MCK2) through one or more unknown cellular receptors while infection of fibroblast occurs independent of MCK2 and is mediated by cell-expressed neuropilin 1. Applying a CRISPR screen, we now identified that the MHC-Ia/Beta-2-microglobulin (B2m) complex serves as an entry port for MCK2-mediated infection of macrophages. Further analyses revealed that MCK2-dependent infection requires expression of the MHC-Ia haplotypes H-2b and H-2d but not H-2k. The importance of the MCK2-MHC-I-pathway for primary infection and viral dissemination was highlighted by experiments with B2m-deficient mice, which lack surface expression of MHC-I molecules. In those mice, intranasally administered MCK2-proficient MCMV could not infect alveolar macrophages and subsequently failed to disseminate into the salivary glands. The identified molecular pathway used by MCMV to infect lung resident macrophages provides essential knowledge for understanding cytomegalovirus-induced pathogenesis, tissue targeting, and virus dissemination.

## Introduction

Human cytomegalovirus (HCMV) is a double-stranded DNA β-herpesvirus that infects approximately 83% of the worldwide population (Zuhair et al., 2019). After contact with body fluids from a virus-shedding individual (Cannon et al., 2011), HCMV usually quickly spreads to multiple organs, most likely carried within abortively infected monocytes (Sinzger and Jahn, 1996; Smith et al., 2004). Nevertheless, in the immunocompetent host, HCMV infection is usually asymptomatic due to efficient virus control by natural killer (NK) cells and T cells (Klenerman and Oxenius, 2016). However, the virus manages to establish latency in endothelial cells, CD34^+^ bone marrow progenitor cells and myeloid cells, allowing virus persistence for a lifetime of the host (Jarvis and Nelson, 2002; Khaiboullina et al., 2004; Mendelson et al., 1996). During latency, viral reactivation is actively suppressed by humoral and cellular immune responses (Picarda and Benedict, 2018; Poole and Sinclair, 2015). Therefore, especially endangered from HCMV infection or reactivation are persons with immature or compromised immune systems, including newborns and transplant patients (Griffiths and Reeves, 2021). In these vulnerable populations, HCMV induces organ-specific manifestations of inflammation such as interstitial pneumonitis and hepatitis.

HCMV is strictly adapted to human cells but the murine cytomegalovirus (MCMV) is a reliable mouse model for studying patho-mechanisms of CMV infection and strategies for intervention (Fisher and Lloyd, 2021; Reddehase and Lemmermann, 2018). The course of MCMV infection in mice depends on the route of infection, that also model certain aspects of the human disease caused by HCMV (Fisher and Lloyd, 2021). The intranasal route of MCMV administration has proven to be an excellent model (Oduro et al., 2016) to mimic the potential natural site of primary infection in mice (Farrell et al., 2016a) and also resembles likely HCMV entry through oral/nasal mucosa in humans (Mayer et al., 2017; Wejse et al., 2001). Upon intranasal administration, MCMV infects lung airway epithelial cells type 2 (AEC2) and macrophages (Stahl et al., 2013, 2015) and induces robust immune responses in the lungs (Lueder et al., 2018). Importantly, MCMV entry into macrophages and viral dissemination from the lungs is dependent on MCMV-encoded chemokine 2 (MCK2) (Ma et al., 2021; Stahl et al., 2015), a CC-chemokine-like product of spliced ORF m131/129 (MacDonald et al., 1999). It has been reported that MCK2 attracts monocytes to the site of infection through CX3CR1 signaling (Daley-Bauer et al., 2014) and enables the virus to disseminate by abortively infected myeloid cells, including monocytes, macrophages, and/or dendritic cells (Baasch et al., 2020; Farrell et al., 2017; Zhang et al., 2021). Upon intranasal infection, MCMV disseminates primarily to salivary glands (Oduro et al., 2016; Stahl et al., 2015), an organ that allows viral persistence and shedding for transmission to new hosts. Little is known about host factors involved in MCMV cell entry, impeding our understanding of primary infection and virus dissemination. Recently, Lane et al. suggested that neuropilin 1 (Nrp1) is the sole cellular receptor for MCMV infection of a broad range of cell types (Lane et al., 2020). In contrast, HCMV uses divergent cellular receptors, depending on the target cell type. To infect fibroblasts, HCMV uses a trimer consisting of glycoproteins L (gL) / gH / gO to bind to platelet-derived growth factor receptor alpha (PDGFRα) (Kabanova et al., 2016; Soroceanu et al., 2008; Stegmann et al., 2017), although several other interactors have also been detected (Martinez-Martin et al., 2018). On the other hand, the gH/gL/UL128/UL130/UL131 pentamer mediates HCMV entry into myeloid, endothelial, and epithelial cells (Gerna et al., 2005; Hahn et al., 2004). Recently, the HCMV pentamer complex was shown to bind several host candidates, including Neuropilin 2, CD147, OR14I1, and CD46 (Martinez-Martin et al., 2018; Stein et al., 2019; Vanarsdall et al., 2018; Xiaofei et al., 2019). Similarly to HCMV (Nguyen and Kamil, 2018), MCMV also has two envelope protein complexes that mediate virus tropism: a gL/gH/gO trimer and an alternative entry complex in which MCK2 replaces gO (Scrivano et al., 2010; Wagner et al., 2013). In contrast to fibroblasts, MCMV infection of macrophages strictly depends on MCK2 (Pontejo and Murphy, 2019; Stahl et al., 2015; Wagner et al., 2013), indicating the existence of additional cellular receptors that allow MCMV to infect cells.

In the present study, we applied a genome-wide CRISPR/Cas9 screening and identified major histocompatibility class I (MHC-I) molecules to serve as host receptors for MCK2-dependent MCMV entry into cells. Moreover, H-2^d^ and H-2^b^ alleles were crucial for viral entry into the macrophages *in vivo* and for virus dissemination from the lungs into the salivary glands.

## Results

### The MCK2 protein is crucial for MCMV macrophage infection

To establish an *in vitro* model for detection of cellular receptors for viral MCK2, we initially compared infectivity of a Smith strain-derived, BAC-cloned MCMV strain with defective MCK2 (MCMV-3D) to an MCMV strain in which the MCK2 locus had been repaired, so that the virus expressed MCK2 (MCMV-3DR) (Stahl et al 2015). We used those viruses to infect NIH/3T3 cells, a mouse embryonic fibroblast cell line, and Raw 264.7, a monocyte/macrophage-like cell line. As previously described, both MCMV-3D and MCMV-3DR carry several reporters, including the red fluorophore mCherry and the enzyme Gaussia luciferase, that can be used to analyze infection efficiencies and the ovalbumin-derived peptide SIINFEKL to study cognate activation of transgenic MHC class-I restricted activation OT-I T cells (Marquardt et al., 2011). Using flow cytometry, we determined viral infectivity at different multiplicity of infection (MOI) as a percentage of mCherry expressing cells 20-24h after infection before the virus starts to spread from cell to cell (Figure 1A). Quantitative analysis revealed that both MCMV-3D and -3DR infected equally well NIH/3T3 fibroblasts (Figure 1B). In contrast, RAW 264.7 cells were 10x more susceptible to MCMV-3DR than to MCMV-3D infection (Figure 1B).

**Figure 1.**
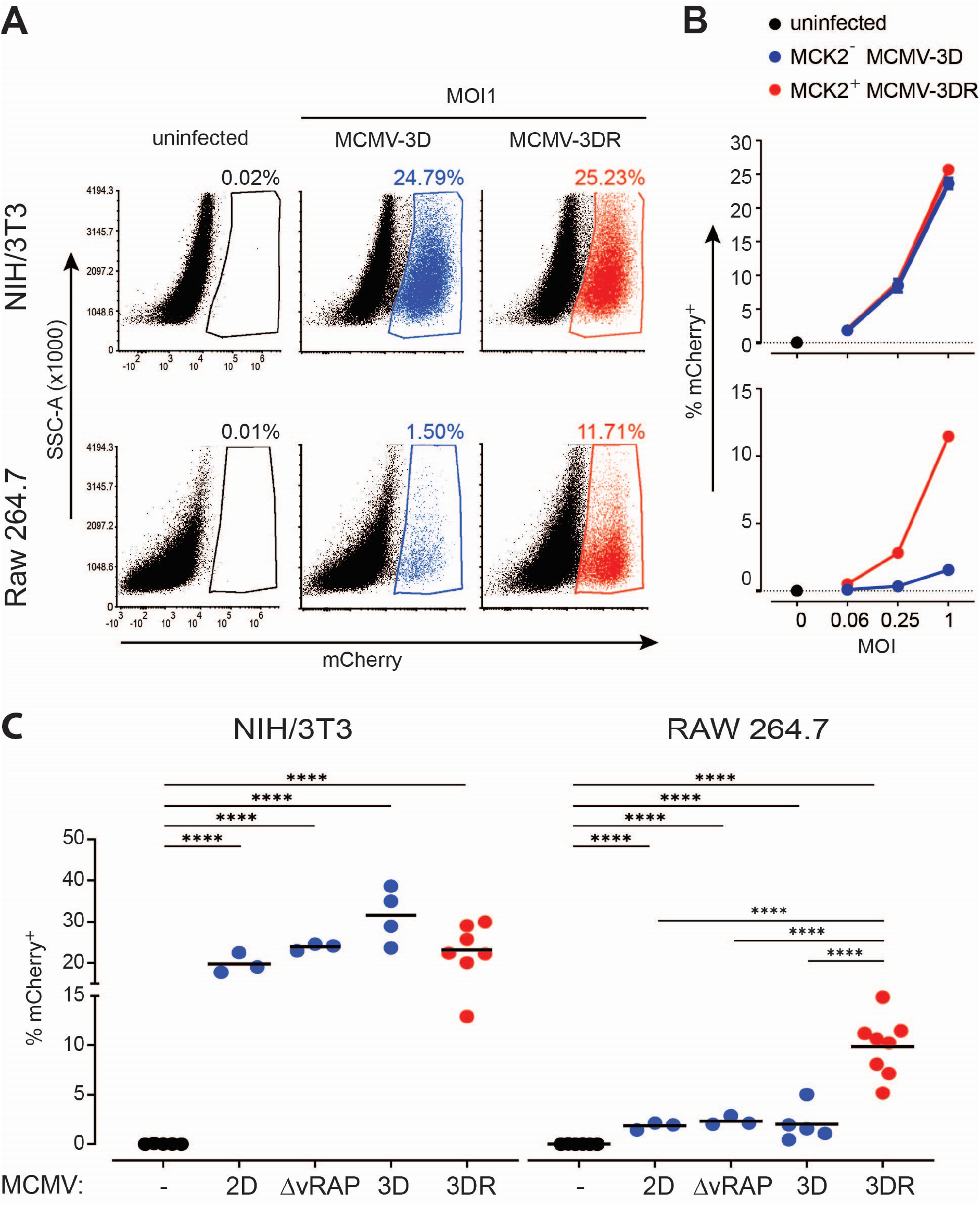
MCK2 enables MCMV to infect macrophages. **(A)** Representative flow cytometry dot plots of uninfected 3T3/NIH fibroblasts and Raw 264.7 monocyte/macrophage cells and cells at 20 h post infection (p.i.) with an MCMV strain devoid of MCK2 (MCMV-3D) compared to strain that expresses MCK2 (MCMV-3DR). **(B)** Quantification of infected cells measured as percentage of mCherry positive NIH/3T3 fibroblasts or Raw 264.7 monocytes/macrophages by flow cytometry at 20 h p.i. with indicated doses of MCMV-3D or MCMV-3DR. **(C)** Infectivity of non-MCK2-expressing (MCMV-2D, MCMV-3DΔVrap; MCMV-3D) and MCK2-expressing (MCMV-3DR) MCMV strains on NIH/3T3 fibroblasts and RAW 264.7 monocytes/macrophages. **(A-C)** All infections were performed at multiplicity of infection (MOI) of 1 and expression of mCherry protein encoded by the virus was measured with flow cytometry at 20 h p. i.. Data are from 2-7 independent experiments. Dots represent the mean of technical triplicates, error bars are SEM, line is the mean value per group. Statistical analysis: Welch’s ANOVA test followed by Dunnett’s T3 multiple comparison test; ****p < 0.0001.

To further validate these results, we used our bone marrow-derived, immortalized Cas9-Hoxb8 hematopoietic progenitor cell line that was differentiated into macrophages in the presence of recombinant mouse macrophage colony-stimulating factor (M-CSF) (Hammerschmidt et al., 2018). As expected, on day 9 of *in vitro* differentiation, the progenitors uniformly expressed the macrophage markers CD11b, F4/80, CD64, and partially CD115 (Figure S1A). In line with the data obtained from our experiments on the RAW 264.7 cells, Cas9-Hoxb8 derived macrophages were also more permissive to infection with the MCK2-proficient MCMV-3DR than to MCK2-deficient MCMV-3D (Figure S1B,C).

Additionally, we performed parallel infections of RAW 264.7 macrophages and NIH/3T3 fibroblasts with two additional MCK2-deficient MCMV variants, named MCMV-2D (lacking the SIINFEKL peptide) and MCMV-3D-ΔvRAP (deficient for the viral repressors of antigen presentation encoded by m06 and m152) (Halle et al., 2016). Analysis of mCherry expression by flow cytometry revealed that all tested virus strains infected NIH/3T3 fibroblasts at similar levels (Figure 1C) while all MCMV strains lacking MCK2 were 4.2 -5.3 times less efficient than the MCK2 expressing MCMV-3DR in infecting macrophages (Figure 1C). These data were also validated by determining Gaussia luciferase released from infected cells in the cell culture supernatant (Figure S2). Altogether, these data confirmed our previous *in vivo* observations that the MCMV-encoded chemokine MCK2 is required for macrophage infection.

### *MCK2*^*+*^ *MCMV enters into macrophages independently of* Nrp1 *or* CX3CR1

To examine whether Nrp1, a recently described receptor for the K181 strain of MCMV (Lane et al., 2020), mediates MCK-2-dependent MCMV entry into cells, we knocked out *Nrp1* on NIH/3T3 and RAW 264.7 macrophages using CRISPR/Cas9 (Figures 2A and S3). As a control, we nucleofected the cells with CRISPR/Cas9 ribonucleoparticles (RNPs) containing a crRNA without specificity for the mouse, rat, or human genomes (Figure 2A). Those control cells showed the same levels of infectivity with MCMV-3D or -3DR strains as their non-nucleofected counterparts (Figure S4). In contrast, the entry of MCMV-3D was almost completely blocked in *Nrp1*^*-/-*^ NIH/3T3 fibroblasts (Figure 2B). This result confirms that Nrp1, despite its low expression levels (Figure 2A), is the main receptor for MCK2-deficient MCMV strains on NIH/3T3 fibroblasts (Lane et al., 2020). However, MCMV-3DR infection was only reduced by 28.7% ± 23.7% in *Nrp1-*deficient fibroblasts (mean ± SD; Figure 2B). Similarly, lack of *Nrp1* had no significant impact on MCK2-dependant infection of Raw 264.7 cells at the low MOI of 1 (Figure 2B) and was only detectable at a high MOI of 5 (Figure S5). Together, these data suggest the existence of additional receptor(s) that allow MCK2-mediated MCMV entry into macrophages.

**Figure 2.**
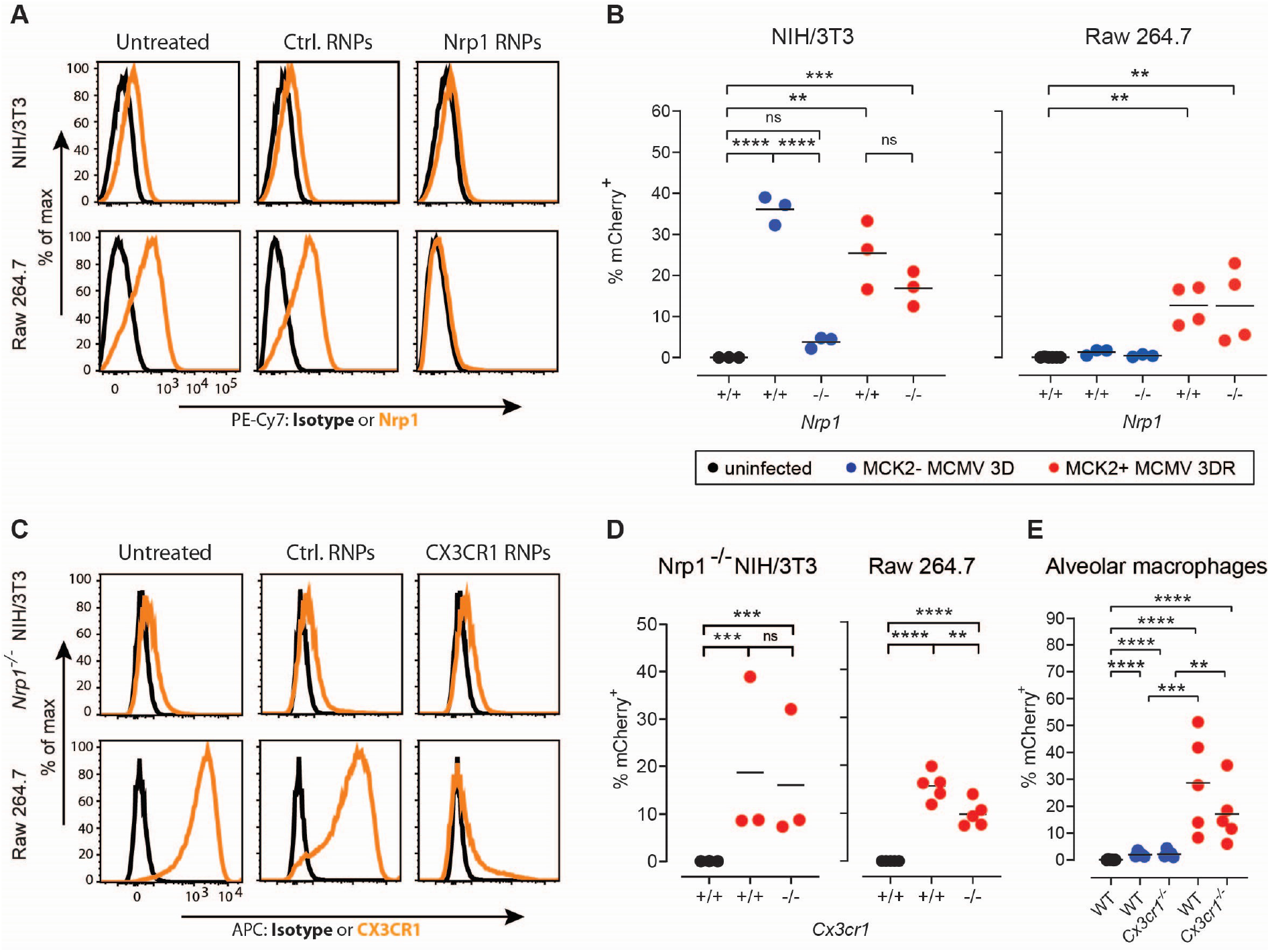
Neuropilin-1 (Nrp1) and CX3CR1 do not mediate MCK2-dependent MCMV infection of macrophages. **(A)** Representative flow cytometry histograms plots of Nrp1 levels on NIH/3T3 fibroblasts or RAW 264.7 monocyte/macrophage cells that were nucleofected with control CRISPR/Cas9 ribonucleoparticles (Ctrl. RNPs) or CRIPSR/Cas9 ribonucleoparticles targeting neuropilin 1 (Nrp1 RNPs). **(B)** Quantification of mCherry signal at 20 h p. i. with indicated MCMV strains from cells treated with Ctrl. RNPs or Nrp1 RNPs. **(C)** Representative flow cytometry histograms plots of CX3CR1 levels on NIH/3T3 fibroblasts or RAW 264.7 monocyte/macrophage cells that were nucleofected with Ctrl. or CX3CR1 RNPs. **(D)** Quantification of mCherry signal at 20 h p. i. with MCMV-3DR from cells treated with Ctrl. RNPs or CX3CR1 RNPs. **(E)** Quantification of mCherry signal at 20 h p. i. with MCMV-3D or -3DR from alveolar macrophages collected by broncho-alveolar lavage from wild-type (WT) or *Cx3cr1*^*-/-*^ mice. **(B**,**D)** Data are from 3-4 independent experiments. One dot equals a mean of the triplicates from one experiment, line at mean value per group. (E) Data are from 2 experiments with 5-6 animals per group (dots) and line represents mean value per group. (C,D,E) Statistical analysis: One-way ANOVA test followed by Sidak’s multiple comparison test; ns, not significant; ** p < 0.01, *** p < 0.001, ****p < 0.0001.

As MCK2 has sequence homology to CC chemokines (MacDonald et al., 1997, 1999), chemokine receptors are the most likely candidates for MCK2 mediated cell infection. However, we have previously shown that MCMV-3DR can infect alveolar macrophages deficient for either CCR1-, CCR2-, CCR5-, CCR7-, or CCR9 (Stahl et al., 2015). On the other hand, MCK2 was reported to mediate the recruitment of patrolling monocytes through the chemokine receptor CX3CR1 (Daley-Bauer et al., 2014). We therefore used CRISPR/Cas9 RNPs to knockout this receptor on *Nrp1*^*-/-*^ NIH/3T3 and *Nrp1*^*+/+*^ Raw 264.7 cells (Figure 2C and S3) and analyzed their susceptibility towards infection with MCK2-proficient MCMV. As CX3CR1 was almost undetectable on NIH/3T3 fibroblasts (Figure 2C), it was not surprising that MCMV-3DR infected at similar extents *Cx3cr1*^*-/-*^ and *Cx3cr1*^*+/+*^ NIH/3T3 cells (Figure 2D).

On the other hand, *Cx3cr1*^*-/-*^ Raw 264.7 cells were on average 33 % less efficiently infected than their *Cx3cr1*^*+/+*^ counterparts (Figure 2D). To validate the role of CX3CR1 in MCK2-mediated MCMV infection of alveolar macrophages, we next collected broncho-alveolar lavage (BAL) cells from untreated mice, in which alveolar macrophages represent the dominant (>85%) cell population (Figure S6A) (Bošnjak et al., 2021; Stahl et al., 2015). In line with previous reports (Gibbings et al., 2017), alveolar macrophages did not express CX3CR1 (Figure S6B). For both virus strains no significant differences could be observed regarding their infectivity of alveolar macrophages derived from wild type or *Cx3cr1*^*-/-*^ mice (Figure 2E). Together, these data indicate that CX3CR1 at most, only partially mediates MCMV entry into macrophages and further support the existence of additional receptors important for MCMV infection.

### Genome-wide CRISPR screen identifies the MHC-I/β2-microglobulin complex and CD81 as target structures for MCK2-mediated MCMV entry

To identify receptors that mediate cell susceptibility towards MCK2 expressing MCMV, we performed a genome-wide CRISPR screen using a mouse CRISPR knockout pooled library (Brie; Addgene). This library targets 19,674 mouse genes, each with 4 single-guide RNAs (sgRNA) (Doench et al., 2016; Sanson et al., 2018). The screen was designed to enrich for cells resistant to virus infection by sorting out mCherry-negative cells from library-containing cells infected with MCMV-3DR at an MOI that was close to infecting all cells in the assay. For the screen, we used *Nrp1*^*-/-*^ NIH/3T3 fibroblasts which required a very high MOI of 16 to infect 75% of cells (Figure S7). Those cells were initially transduced to stably express a functional Cas9-protein before they were infected with library-containing lentiviral vectors at an MOI of 0.33 to limit the delivery of sgRNAs to one per cell. Puromycin-selected library-containing *Nrp1*^*-/-*^ NIH/3T3 fibroblasts were then infected with MCMV-3DR at an MOI 10 and surviving mCherry^-^ cells were sorted 20 h p.i. After five hours in culture, these cells were again infected with the same MOI, reducing the starting population to approximately 7% of cells that remained mCherry-negative. We expanded these cells for 2 days, allowing them to divide twice before we performed a third MCMV-3DR infection with an MOI 16. One set of sorted mCherry-negative cells was then used for analysis, while the other set of mCherry-negative cells was infected for a fourth time with an MOI of 16. From sorted mCherry-negative cells resistant to MCMV infection, we extracted genomic DNA and sequenced PCR-amplified gRNA encoding regions to identify sgRNAs that were enriched in cells resistant to MCMV-3DR infection compared to the untreated input population (Figure 3A). Bioinformatic analysis identified overrepresented sgRNAs in the terminally sorted mCherry-negative cell populations (Figure 3B). To reduce the possibility of false-positive results, we overlapped the lists of sgRNAs generated in each biological replicate, revealing six genes potentially implicated in MCK2-mediated MCMV entry into the cells (Figure 3C). Three of these genes, *B2m, Tapbp*, and *Tap1*, are crucial for MHC-I expression on the cell surface (Raghavan et al., 2008). Of the remaining hits, only *Cd81* encodes for an extracellular protein, a tetraspanin that was implicated in influenza and hepatitis-C-virus entry in human cells (Florin and Lang, 2018). Protein disulfide isomerase associated 3 is a chaperone protein encoded by *Pdia3* that prevents protein aggregation and is upregulated upon influenza inflammation in lung cells (Chamberlain et al., 2019). Finally, *Keap1* encodes for an inhibitor of the nuclear factor erythroid 2-related factor 2 (Nrf2), a transcription factor important for the suppression of oxidative stress induced by viral infection (Wang et al., 2022). Altogether, these data implied that the cell-surface expressed MHC-I/B2m complex and CD81 serve for MCK2-mediated MCMV entry into the cells.

**Figure 3.**
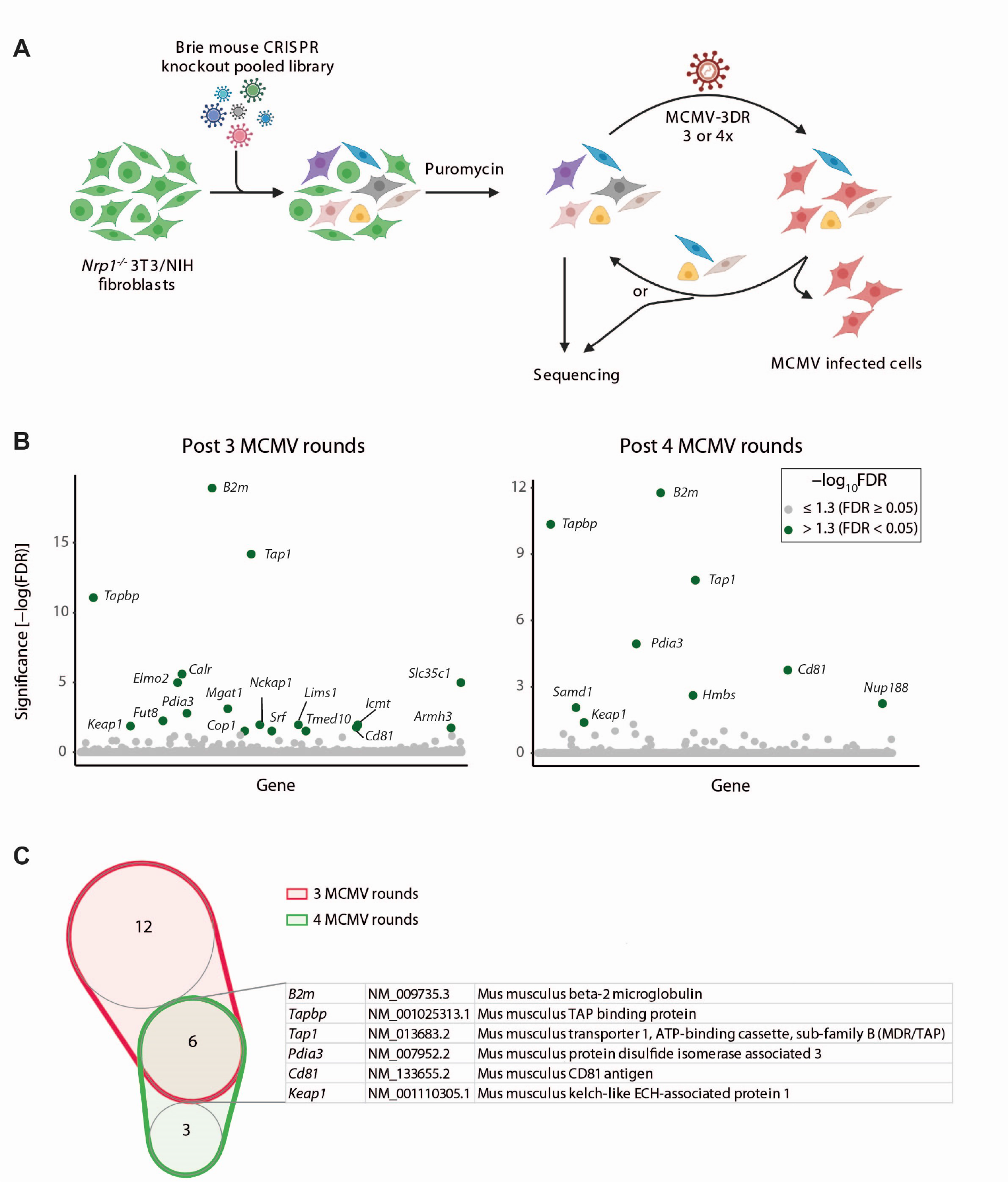
Genome wide CRISPR screen identified CD81 and B2m/MHC-I complex as putative factors for MCMV entry. **(A)** Scheme of the CRISPR screen protocol created with BioRender.com. **(B)** Screen results on the *Nrp1*^*-/-*^ NIH/3T3 fibroblasts after the indicated number of MCMV-3DR infection rounds. Vertical axis indicates significance of sequence enrichment in surviving cells compared to input cell population for all genes arbitrarily spread along the horizontal axis. **(C)** List of genes shared between the two screens.

### Functional validation of MHC-I as a receptor for MCK2-mediated MCMV entry into macrophages

To validate the targets from the CRISPR screen, we knocked out *B2m* and *Cd81* in *Nrp1*^*-/-*^ NIH/3T3 fibroblasts and RAW 264.7 monocytes/macrophages. In cells treated with CD81 Cas9 RNPs, CD81 could not be detected on the cell surface (Figure 4A). As MHC-I expression is dependent on beta-2-microglobulin (Li et al., 2016), MHC-I protein was almost undetectable on the surface of both NIH/3T3 (H-2^q^ haplotype) and Raw 264.7 (H-2^d^ haplotype) cells treated with RNPs targeting *B2m* (Figure 4B). Upon exposure to the herpesviral particles, we observed that both *B2m*^*-/-*^*Nrp1*^*-/-*^ NIH/3T3 and *B2m*^*-/-*^ Raw 264.7 cells were almost completely protected from MCMV-3DR infection (Figure 4C and 4D). In contrast, cells carrying mutations in the *Cd81* gene were only partially protected and the percentage of mCherry^+^ *Cd81*^*-/-*^*Nrp1*^*-/-*^ NIH/3T3 and *Cd81*^*-/-*^ Raw 264.7 cells was reduced by 36%±2% and 48%±12%, respectively (Figure 4C and 4D).

**Figure 4.**
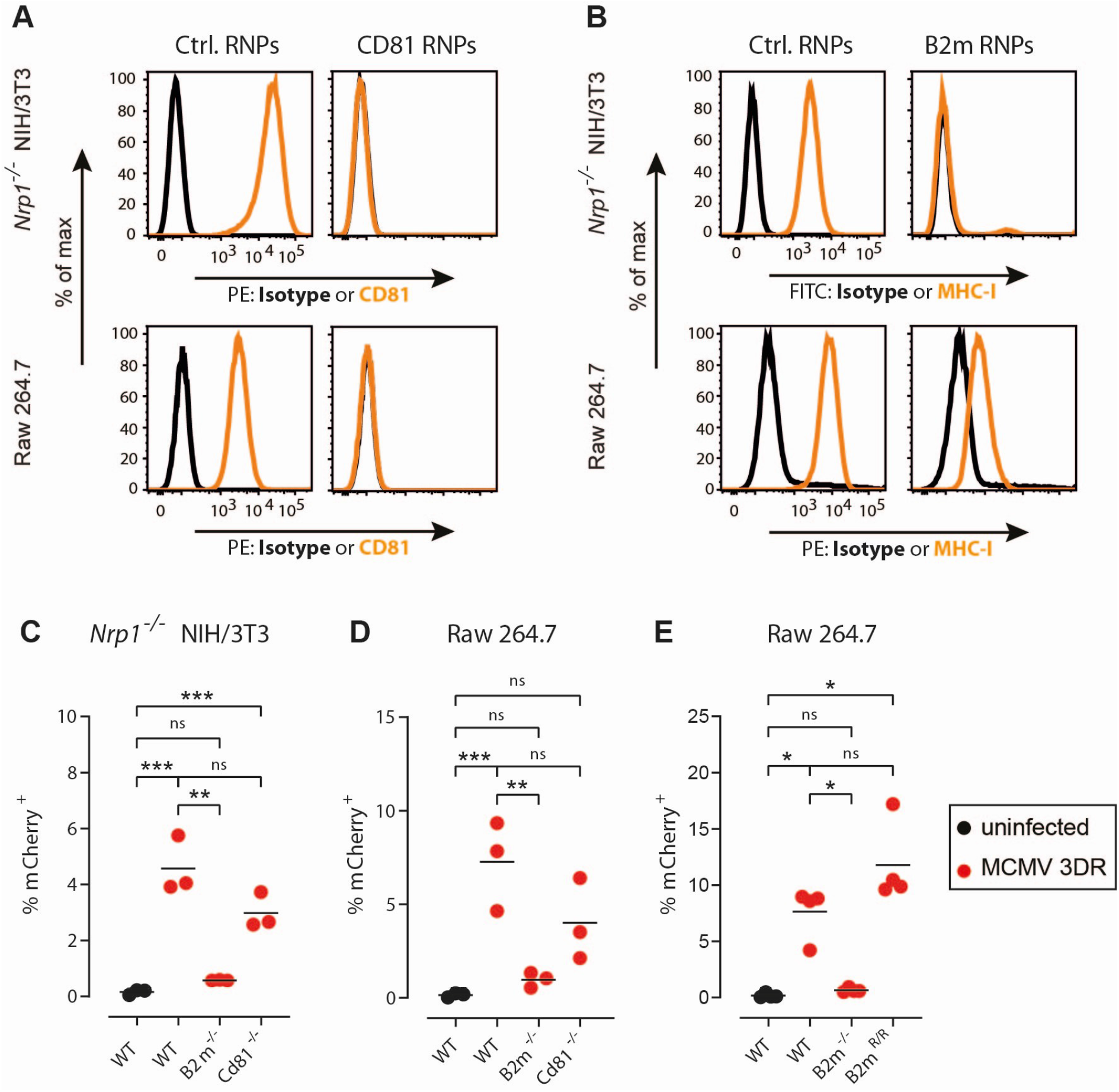
The B2m/MHC-I complex rather than CD81 mediates MCMV entry into macrophages. **(A**,**B)** Representative flow cytometry histograms depicting (A) CD81 or (B) MHC-I expression on *Nrp1*^*-/-*^ NIH/3T3 fibroblasts and RAW 264.7 macrophages nucleofected with control CRISPR/Cas9 ribonucleoparticles (Ctrl. RNPs) or CRIPSR/Cas9 ribonucleoparticles targeting (A) CD81 (CD81 RNPs) or (B) Beta-2 microglobulin (B2m RNPs). **(C**,**D)** Quantification of mCherry signal at 20 h p. i. with MCMV-3DR from (C) *Nrp1*^*-/-*^ NIH/3T3 fibroblasts and (D) RAW 264.7 macrophages cells knockout for indicated genes. **(E)** Quantification of mCherry signal at 20 h p. i. with MCMV-3DR from *B2m*^*-/-*^ Raw 264.7 cells in which single-nucleotide mutation within *B2m* gene was repaired using CRISPR/Cas9-mediated gene editing (*B2m*^*R/R*^ cells; Figure S7) restored susceptibility to MCMV-3DR infection. **(C**,**D**,**E)** Data are from 3-4 independent experiments. One dot equals a mean of the triplicates from one experiment, line at mean value per group. Statistical analysis: One-way ANOVA test followed by Sidak’s multiple comparison test; ns, not significant; ** p < 0.01, *** p < 0.001, ****p < 0.0001.

To validate that observed differences in MCK2-mediated infectivity are indeed related to mutations within the *B2m* and not by off-target genome editing effects, we created cells with a corrected mutation within the *B2m* gene using CRISPR/Cas9-mediated homology-directed repair (*B2m*^*R/R*^ cells; Figure S8). Importantly, these *B2m*^*R/R*^ cells became again fully susceptible to MCMV-3DR infection (Figure 4E), confirming the importance of MHC-I/B2m complex in MCK2-mediated MCMV entry into the cells.

To validate further our results, we next collected BAL cells from WT and *B2m*^*-/-*^ mice and infected them *in vitro* with MCMV-3D or -3DR. In line with our results on Raw 264.7 cells, we found that MCMV-3DR was unable to infect BAL cells isolated from *B2m*^*-/-*^ mice (Figure 5A). Moreover, BAL cells isolated from MCMV-susceptible C57BL/6 (H-2^b^ haplotype) and BALB/c mice (H-2^d^ haplotype) but not MCMV-resistant C3H mice (H-2^k^ haplotype) showed higher infection rates toward MCMV-3DR than MCMV-3D virus (Figure 5B). These data, therefore, not only confirmed the importance of the MHC-I/B2m complex in MCK2-mediated MCMV entry but also indicated that the MHC-I molecule, rather than B2m, serves as a binding partner for viral MCK2.

**Figure 5.**
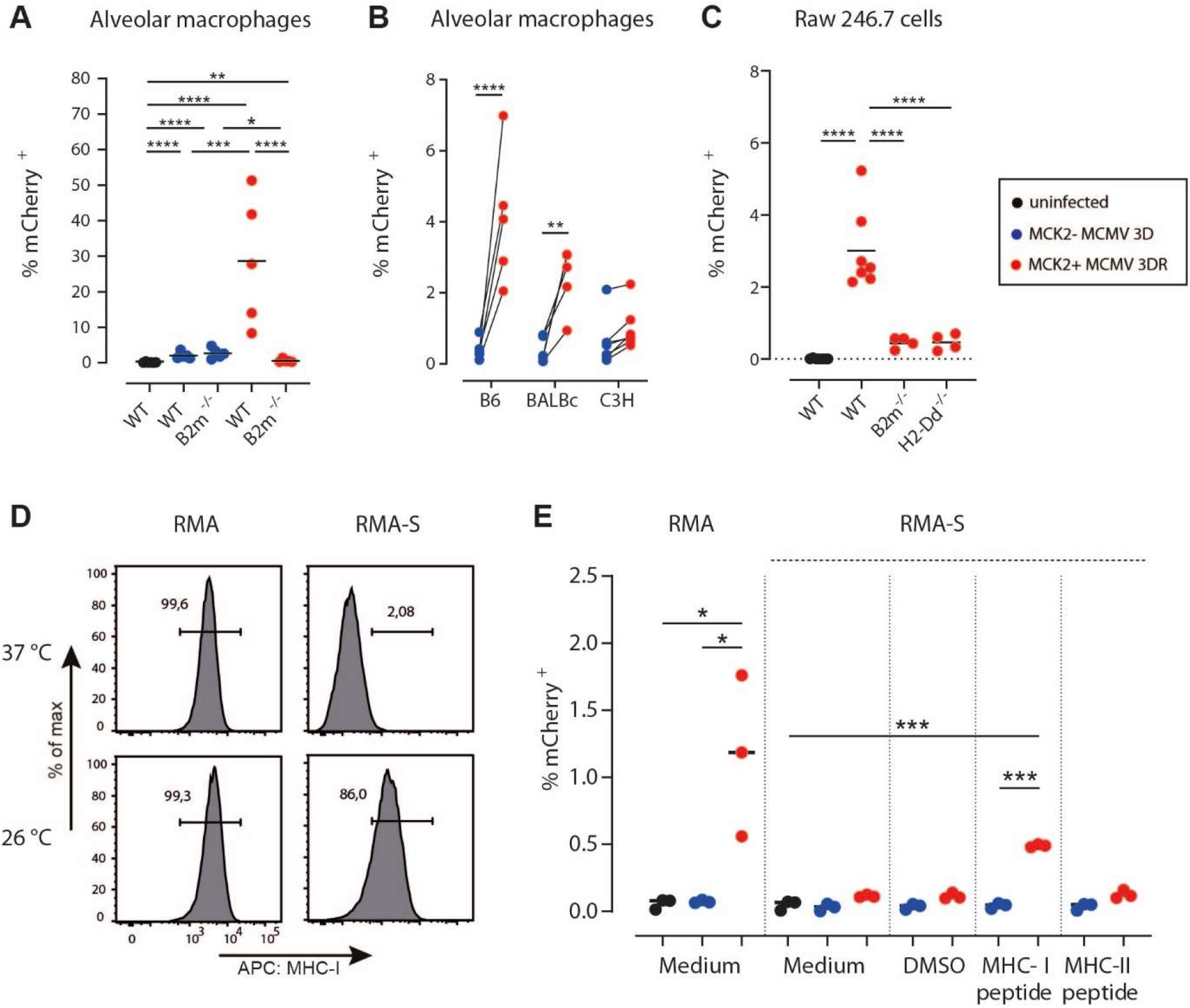
MCK-2 mediated MCMV-3DR entry depends on the MHC-I. **(A)** Percentage of mCherry^+^ broncho-alveolar lavage cells collected from wild type (WT) B6 or B2m^-/-^ mice at 20 h p.i. with MCK2-deficient MCMV-3D or MCK2-proficeint MCMV-3DRat MOI 1. Pooled data from two experiments with 5-6 mice per group. **(B)** Broncho-alveolar lavage (BAL) cells collected from wild type (WT) B6, BALB/c, or C3H mice were split into two halves and then infected *in vitro* with MCK2-deficient MCMV-3D or MCK2-proficeint MCMV-3DR at MOI 1. Data indicate percentage of mCherry^+^ cells at 20 hr p.i. and are pooled from 2 experiments with 4-6 mice per group. **(C)** Quantification of mCherry signal in WT, *B2m*^*-/-*^, and *H-2Dd*^*-/-*^ Raw 264.7 cells at 20 hours p. i. with MCMV-3DR at MOI 1. Data are from 4-7 independent experiments. **(D)** Expression of MHC-I on RMA and RMA-S cells grown at 37°C (upper row) and then kept for 26-28 hours at 26°C (lower row). **(E)** RMA and RMA-S cells were kept for 26-28 h at 26°C before medium, medium with DMSO, medium with MHC-I binding SIINFEKL peptide, or MHC-II binding peptide were added as indicated. After additional 4 h of incubation, cells were infected with MCMV-3D or -3DR at MOI 2. After infection, the cells were further incubated at 26°C for 20-22 h, when percentage of mCherry^+^ cells was analyzed by flow cytometry. Data are from 3 independent experiments. **(A**,**B**,**C**,**E)** One dot equals the mean of the technical duplicates or triplicates, line the mean value per group. Statistical analysis: Welch’s ANOVA test followed by Dunnett’s multiple comparison test (RMA cells) or one-way ANOVA test followed by Sidak’s multiple comparison test (all other cells); * p < 0.05, ** p < 0.01, *** p < 0.001, ****p < 0.0001.

To confirm this hypothesis, we next created *H2-D1*^*-/-*^ Raw 264.7 cell line using CRISPR/Cas9 (Figure S9) and found that knocking out only this MHC-I allele decreased the susceptibility of cells towards MCMV-3DR to the level of *B2m*^*-/-*^ Raw 264.7 cells (Figure 5C). In addition, we also used the lymphoma cell lines RMA and RMA-S. In contrast to RMA cells, RMA-S carry a mutation in the TAP-2 gene, which impairs peptide loading into MHC-I molecules and their transport to the cell surface. However, incubation of cells at 24°C to 28°C increases the expression of MHC-I molecules (Figure 5D), which can be further stabilized by loading them with appropriate peptides (Schumacher et al., 1990). It has been shown that treatment of RMA-S cells with MHC-I-binding peptides at 28°C makes these cells susceptible to MCMV infection (Wykes et al., 1993). In line with those results, MCMV-3DR was unable to infect RMA-S cells at 26°C unless they were loaded with a peptide that can stabilize H-2K^b^ molecules (Figure 5E). Importantly, MCMV-3D could not infect either RMA or RMA-S cells (Figure 5E). Altogether, these data confirm that properly folded MHC class Ia molecules, primarily H-2^d^ and H-2^b^ alleles, serve as an entry receptor to which MCMV binds via MCK-2.

### B2m^-/-^ mice are protected from MCMV infection in vivo and virus dissemination to the salivary gland

To validate our *in vitro* findings on the course of MCMV infection in mice, we intranasally infected age-matched cohorts of wild-type B6 (WT) and *B2m*^*-/-*^ mice with 1×10^6^ plaque-forming units (PFU) of MCMV-3D or MCMV-3DR. In line with our previous observations (Stahl et al., 2013, 2015), we found that at 1 day post infection (dpi) MCMV-3DR infected both CD11c^+^ cells positioned mainly in alveolar space and CD11c^-^ cells positioned in lung epithelium of WT mice (Figure 6A,B). In sharp contrast, in *B2m*^*-/-*^ mice mCherry signals encoded by MCMV-3DR were almost exclusively found in CD11c^-^ cells, resembling the pattern of MCMV-3D infection that was mainly restricted to CD11c^-^ stroma cells in both WT and *B2m*^*-/-*^ mice (Figure 6A,B). Analysis of lung cells with flow cytometry further confirmed that the MCMV-3DR infected mCherry^+^CD11c^+^ cells in WT mice were indeed alveolar macrophages, defined as CD11c^+^CD11b^low^SiglecF^+^CD103^-^ cells (Figure 6C,D and S10). In line with histological observations, quantification of flow cytometry data indicated that alveolar macrophages from *B2m*^*-/-*^ mice were resistant to MCMV-3DR infection (Figure 6E). In agreement with our previous findings (Stahl et al., 2015), MCMV-3D could not infect alveolar macrophages in either of the two mouse strains used (Figure 6E). Not surprisingly, we also found significantly higher luciferase activity in the lung homogenates of WT compared to *B2m*^*-/-*^ mice infected with MCMV-3DR (Figure 6F), suggesting that the MCK2-MHC-I mediated entry pathway enables MCMV to establish a foothold in lungs by infecting alveolar macrophages after intranasal infection.

**Figure 6.**
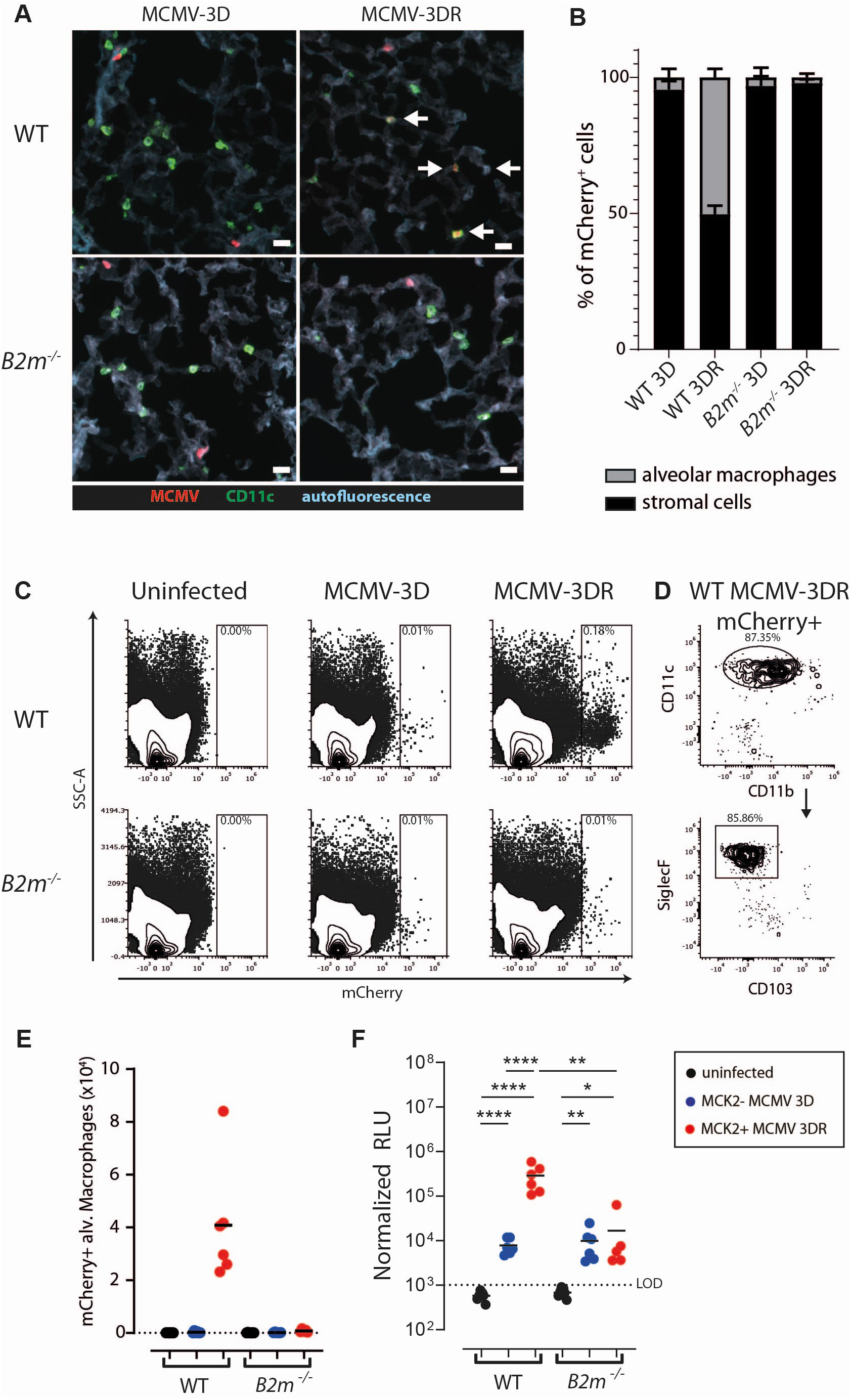
Alveolar macrophages in B2m^-/-^ mice are protected from in vivo MCMV-3DR infection. **(A)** Representative photomicrographs of anti-CD11c stained lung sections indicating mCherry^+^ (infected) cells in the lungs of wild-type B6 (WT) or *B2m*^*-/-*^ mice at 1 day post infection (dpi) with MCMV-3D or -3DR (dpi). Arrows indicate CD11c^+^mCherry^+^ cells. Bar = 20 µm. **(B)** Frequencies of infected alveolar macrophages (CD11c^+^mCherry^+^ positioned in the alveolar space) and stroma cells (CD11c^-^mCherry^+^ positioned in lung epithelium) were quantified. One section / mouse was analyzed with a minimum of 15 mCherry^+^ cells/mouse analyzed. **(C)** Representative contour plots indicating mCherry signal within all live lung cells of uninfected mice or mice at 1 dpi with viruses as indicated. **(D)** Flow cytometric characterization of mCherry^+^ cells in the lungs of WT mice indicated that majority of infected cells are CD11c^+^CD11b^low^SiglecF^+^CD103^-^ alveolar macrophages. **(E)** Number of mCherry^+^ alveolar macrophages, gated as CD11c^+^CD11b^low^SiglecF^+^CD103^-^ cells, quantified from flow cytometry data from the lungs of WT or *B2m*^*-/-*^ mice infected as indicated. **(F)** Luciferase activity in the lungs of WT or *B2m*^*-/-*^ mice 1 dpi with MCMV-3D or -3DR. LOD – limit of detection. **(A-F)** Data were obtained from three independent experiments with n = 5-7 per group. (F) Statistical analysis: Welch’s ANOVA test followed by Dunnett’s T3 multiple comparison test on log_10_-transformed values; * p < 0.05; ** p < 0.01, *** p < 0.001, ****p < 0.0001. RLU values from different experiments were normalized using data from uninfected mice.

Apart from its important role in primary infection, others and we have shown that MCK2 also plays a crucial role in MCMV dissemination to the salivary gland upon virus administration to the lungs (Farrell et al., 2016b; Stahl et al., 2015) or injection into the footpad (Daley-Bauer et al., 2014; Noda et al., 2006). Nevertheless, MCMV titers in the salivary glands of *B2m*^*-/-*^ 129/Sv x C57BL/6 mice two weeks after viral administration into the footpad had been reported not to be different from those detected in *B2m*^*+/-*^ littermates (Polić et al., 1996). As our data indicated the crucial role of MHC-I molecules for MCK2-mediated MCMV entry, we next examined the role of MCK2 in the dissemination to the salivary gland after intranasal administration with MCMV-3D and -3DR in *B2m*^*+/+*^ and *B2m*^*-/-*^ 129/Sv x C57BL/6 mice. At 7 dpi, we found decreased luciferase activity and viral titers in the salivary glands of *B2m*^*-/-*^ mice infected with MCMV-3DR compared to WT mice infected with the same virus (Figure 7A,B). Viral titers detected in salivary glands of *B2m*^*-/-*^ mice infected with MCMV-3DR were comparable to the titers of MCMV-3D infected wt mice, further highlighting the crucial contribution of the MHC-I/MCK2 axis for viral dissemination. In contrast, viral titers in the lungs were comparable among all infected groups at 7 dpi (Figure 7C), although the luciferase activity remained increased in the lung homogenates of WT mice infected with MCMV-3DR (Figure 7D). These data confirmed previous observations that the lack of MCK2 does not impair virus spread within the lungs (Farrell et al., 2016b; Stahl et al., 2015). Moreover, except for low levels of luciferase activity in spleens, no virus-encoded luciferase activity was detected in the livers or kidneys of neither WT nor *B2m*^*-/-*^ mice (Figure S11).

**Figure 7.**
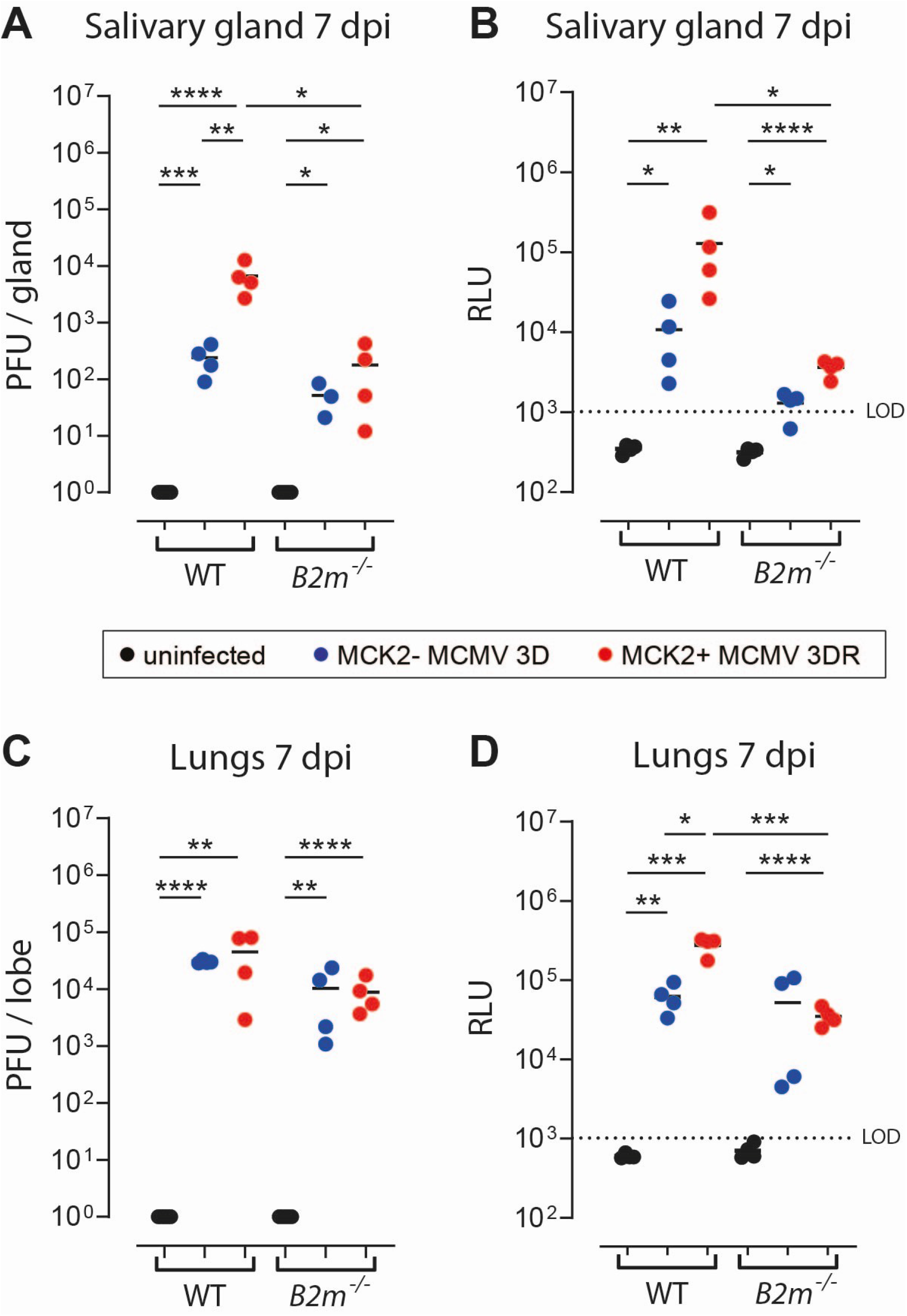
B2m^-/-^ mice are protected from MCMV spread to the salivary gland 7 days after intranasal infection. **(A**,**C)** Virus titers and **(B**,**D)** luciferase activity in **(A**,**B)** salivary glands and **(C**,**D)** lungs of wild-type B6 (WT) or *B2m*^*-/-*^ mice 7 days post infection (dpi) with MCMV-3D or -3DR. LOD – limit of detection. **(A-D)** Pooled data from two experiments with 4 mice per group. Statistical analysis: Welch’s ANOVA test followed by Dunnett’s T3 multiple comparison test on log_10_-transformed RLU or PFU values; * p < 0.05; ** p < 0.01, *** p < 0.001, ****p < 0.0001.

## Discussion

Detection of cellular receptors that serve as entry points for viruses allows insights into the mechanisms required for primary infection, virus spread, and latency within the host. Using a CRISPR screen, we revealed in the present study that MCK2-proficient MCMV infects cells via MHC class Ia molecules.

Here, we focused on the viral entry after respiratory tract infection, where lung AEC2s and alveolar macrophages represent the primary ports of entry into the organism (Farrell et al., 2016b; Stahl et al., 2015). We found that MCK2-proficient MCMV relies primarily on MHC-I to infect alveolar macrophages while the MCK2-deficient virus is restricted to use Nrp1, a recently described as an entry receptor for MCMV (Lane et al., 2020). In the lung, the Nrp1 pathway almost exclusively results in infection of AEC2s. Our finding that infection of the lungs of *B2m*^*-/-*^ mice with MCK2-proficient virus is restricted to stroma cells goes along with this model. These findings suggest that MCMV can use divergent entry pathways to establish a foothold at the primary infection site. Given these differences in virus-cell tropism, it was interesting to see that infection with MCK2-deficient and MCK2-proficient MCMV resulted in comparable viral loads in the lungs of wt and *B2m*^*-/-*^ mice at 7 dpi suggesting that the virus primarily uses the Nrp1 axis to spread within the lung. This finding could potentially be attributed to the high expression levels of Nrp1 in the lungs, where it plays an important role in its organogenesis and function (Joza et al., 2013; Mahmoud et al., 2019; Wild et al., 2012). Although Nrp1 has also been detected on numerous leukocyte populations, including dendritic cells, mast cells, divergent T cell subpopulations, and macrophages (Roy et al., 2017) none of these cell subsets get substantially infected with MCK2-deficient MCMV, suggesting that additional factors are needed to efficiently infect Nrp1 expressing cells. Our data also support this notion since Nrp1 expression levels on NIH/3T3 and Raw 264.7 cells do not correlate with cell infectivity via the Nrp1 axis. Therefore, an important next step would be to determine additional factors such as the binding affinities of viral complexes to their cellular receptors.

If Nrp1 is crucial for viral spread within the lungs, how does the virus profit from MCK2 – MCH-I-mediated infection of alveolar macrophages? Previously, we have shown that depletion of alveolar macrophages in neonates reduces MCMV loads within the lungs, suggesting that these cells contribute to viral replication *in vivo* (Stahl et al., 2015). In contrast, others reported that depletion of alveolar macrophages in adult mice increased MCMV titers in the lung during the acute phase of infection, suggesting that macrophages have a protective role in the immune response to MCMV (Farrell et al., 2016b). These at first glance conflicting data could be explained by a recent study that examined in detail how MCMV infection drastically subverts macrophage identity (Baasch et al., 2021). Interestingly, MCMV infection-induced global transcription re-programming within alveolar macrophages disrupts their ability to phagocyte and secrete inflammatory mediators and increase their migration properties. In contrast, non-infected, by-stander macrophages upregulated type I IFN activation and viral defense response genes, thus contributing to anti-MCMV immune response (Baasch et al., 2021). Therefore, MCK2-proficient virus may infect a higher proportion of neonatal alveolar macrophages, which removal would thus reduce viral loads in the lungs. On the other hand, many alveolar macrophages in adult mice might remain uninfected and contribute to immune protection. Taken together it seems likely that MCK2-mediated infection of alveolar macrophages appears to be an additional viral immunoevasive mechanism.

A broad expression of MHC-I and Nrp1 could explain why MCMV can infect a wide range of cells within its host. However, the susceptibility to the virus seems to depend on the cell type infected. Despite considerably higher levels of Nrp1, Raw cells are less vulnerable to MCK2-deficient MCMV than NIH/3T3 fibroblasts. Similarly, our experiments with RMA cells indicated that these cells required higher MOI for infection with MCK2-proficient MCMV than NIH/3T3 or Raw cells, despite comparable MHC-I expression levels. Together, these data suggest that cell susceptibility to MCMV infection cannot be attributed solely to the protein expression levels of virus receptors. It is possible that differences in receptor recycling or endocytic pathways additionally affect virus internalization, which has to be addressed in future studies.

Besides an important role in primary cell tropism, MCK2 is also essential for MCMV dissemination from the lungs into the salivary glands (Fleming et al., 1999; Jordan et al., 2011; Saederup et al., 2001; Stahl et al., 2015). A large body of evidence indicates that upon intranasal or footpad administration MCMV travels to the salivary glands via blood, hidden in non-productively infected mononuclear cells (Daley-Bauer et al., 2014; Farrell et al., 2017; Noda et al., 2006; Stoddart et al., 1994; Zhang et al., 2021). However, the exact cell type that carries the virus is still under investigation, with evidence that MCMV uses CX3CR1^+^ patrolling monocytes (Daley-Bauer et al., 2014), dendritic cells (Farrell et al., 2017; Ma et al., 2021), myeloid progenitor cells (Noda et al., 2006), or even alveolar macrophages (Baasch et al., 2021) to disseminate to other organs. In addition to cell-mediated dissemination, MCMV can also infect the salivary gland by cell-free viruses in the blood (Zhang et al., 2021). Cell-free dissemination is also presumed to be the main mechanism by which MCK2-deficient MCMV reaches the salivary gland or by which MCK2-proficient virus disseminates in *Ccr2*^*-/-*^ or *Cx3cr1*^*-/-*^ mice that have impaired monocyte migration (Daley-Bauer et al., 2012, 2014; Noda et al., 2006; Stahl et al., 2015). Our finding that MCMV uses at least two different entry pathways for different types of cells identifies a sophisticated molecular framework for virus spread throughout the organism. In *B2m*^*-/-*^ mice the dissemination of MCK2-proficient MCMV from the lungs to the salivary glands was not completely abrogated but reduced to the levels of dissemination observed for MCK2-deficient MCMV in both mouse strains examined. These data suggest that the MCK2 – MCH-I entry pathway serves MCMV to infect mononuclear cells responsible for virus dissemination into the salivary gland. If this axis is affected (by lack of MCK2 or MHC-I), MCMV can still spread to the salivary gland presumably using the cell-free route and infecting the organ via Nrp1-mediated targeting of endothelial cells (Lane et al., 2020). Similarly, the cell-free route of dissemination via Nrp1 could also explain why after subcutaneous or intraperitoneal administration MCK2-deficient and -proficient MCMV strains spread to other organs except for the salivary gland (Fleming et al., 1999; Jordan et al., 2011; Saederup et al., 2001; Stahl et al., 2015). This model can also explain discrepancies between our study and a previous report showing unimpaired MCMV dissemination after footpad administration to salivary glands in *B2m*^*-/-*^ mice (Polić et al., 1996). Although not addressed in that study, it seems not unlikely that the authors used the tissue-culture-grown Smith strain of MCMV carrying a mutation in the MCK2 gene. Alternatively, the virus uses divergent routes to disseminate from the lungs than from the footpad. It has been suggested that the dissemination from the footpad to the salivary glands depends on CX3CR1^+^ patrolling monocytes (Daley-Bauer et al., 2014). As our data indicated that MCK2-proficient virus infects cells also via CX3CR1, MCMV could exploit this chemokine receptor to recruit, enter, and spread via monocytes. However, a later report indicated that after intranasal administration MCMV disseminated independently of CX3CR1 or CCL2 and suggested that the virus uses dendritic cells to spread (Farrell et al., 2019). Therefore, it would be crucial to evaluate the relevance of existing entry pathways for MCMV infection of different cell types, including monocytes and dendritic cells.

Our data also provide a molecular mechanism for why certain mouse strains are resistant to MCMV infection (Chalmer et al., 1977; Selgrade and Osborn, 1974). The present study shows that MCK2-mediated MCMV entry depends on the type of MHC-I allele, as cells expressing H-2^b^ and H-2^d^ alleles were more permissive to infection than cells carrying H-2^k^ alleles. Thus, the resistance of C3H (H-2^k^) mice to MCMV infection seems to be a consequence of the virus’ inability to infect (myeloid) cells and spread within the mice. This hypothesis is in agreement with previous *in vitro* studies indicating the importance of stable expression of correctly folded MHC class I molecules for MCMV infection (Price, 1994; Wykes et al., 1992, 1993).

It has been shown that MCK2 associates with MCMV glycoproteins gH and gL, forming a trimeric gH/gL/MCK2 complex, which is an alternative to the trimeric gH/gL/gO complex (Wagner et al., 2013). Interestingly, the same study showed that virus particles lacking both gO and MCK2 are not infectious unless trans-complemented with either of the proteins. Hence, viruses lacking MCK2 rely exclusively on the trimeric gH/gL/gO complexes to enter cells, suggesting that MCK2^-^ MCMV entry into fibroblasts via Nrp1 is mediated with this trimeric complex. On the other hand, it seems likely that the alternative gH/gL/MCK2 complex is required for MCMV to bind to MHC-I and enter into macrophages. Although this hypothesis requires experimental validation, it also further highlights that MCMV resembles HCMV, which also uses different cellular receptors and viral protein complexes to infect divergent cell types.

Altogether, our data indicate that Nrp1 and MHC-I molecules serve as non-redundant receptors through which MCMV infects various cell types. They provide essential knowledge for understanding cytomegalovirus-induced pathogenesis and virus dissemination.

## Acknowledgments

Alaleh Rezalotfi, Lea Fritz, and Kim Do were supported by the Hannover Biomedical Research School (HBRS) and the Center for Infection Biology (ZIB). We gratefully acknowledge the superb service of Dr. Matthias Ballmaier of the Cell Sorting Core Facility of Hannover Medical School. The CRISPR screen libraries used in this publication were sequenced at the Research Core Unit Genomics at Hannover Medical School. We also thank Ms. Svetlana Piter for providing excellent animal care.

Research in the Förster lab is supported by the German Center for Infection Research TTU 01.938 (grant no 80018019238); by the German Center for Lung Research (grant 82DZL002B1); and by Deutsche Forschungsgemeinschaft (DFG, German Research Foundation) Excellence Strategy EXC 2155 “RESIST” (Project ID39087428), SFB 900/3 (Project B1, 158989968) and FO334/7-1 (all to RF).

## Author contributions

RF and BB conceptualized the work. BB and EH performed CRISPR-screen and analyzed the data, BB, EH, KD, CR, SW, HG performed *in vitro* infection experiments, AR performed CRISPR/Cas9-mediated gene editing and *in vitro* infection experiments, BB, YL, KD, CR, AB performed *in vivo* experiments, YL and KD analyzed FACS data from *in vivo* experiments, AB and YL performed histological staining and analyses, KW and MM prepared virus stocks and determined viral loads in the lungs and salivary glands, MG created Cas9 expressing cell lines, BB and EH designed figure layouts with help from YL, LF and AR. BB, EH, and RF wrote the manuscript. All authors contributed to manuscript draft correction and approved the submitted version.

## STAR Methods

### Resource availability

#### Lead contact

Further information and requests for resources should be directed to and will be fulfilled by the lead contact, Berislav Bošnjak.

#### Materials availability

Cell lines generated by this study are available through request to the lead contact, Berislav Bošnjak with a completed Material Transfer Agreement.

## Materials and methods

### MCMV strains

All MCMV strains used in this study were derived from the Smith strain (pSM3fr) as a bacterial artificial chromosome-cloned MCMV progeny. Virus stocks were produced and titrated in vitro using standard techniques (Halle et al., 2016; Marquardt et al., 2011; Stahl et al., 2015). As described previously, MCMV-2D, MCMV-3D-ΔvRAP, and MCMV-3D carry a frameshift mutation in the m129 region and express truncated and hence dysfunctional MCK2 (Halle et al., 2016; Jordan et al., 2011; Marquardt et al., 2011; Stahl et al., 2015). In MCMV-3DR, the m129 mutation is repaired, so this virus encodes fully functional MCK2 (Halle et al., 2016; Marquardt et al., 2011; Stahl et al., 2015). All strains encode for secretable Gaussia Luciferase and red fluorescent protein mCherry. Additionally, MCMV-3D, MCMV-3DR, and MCMV-3D-ΔvRAP express SIINFEKL peptide. Finally, MCMV-3D-ΔvRAP carries additional mutations in m06/gp48 and m152/gp40, thus lacking two viral regulators of antigen presentation. Of note, experiments for comparative analysis of divergent MCMV strains were performed in parallel with simultaneously titrated virus stocks.

### Cell lines

RAW 264.7 monocytes/macrophages (ATCC® TIB-71™) and NIH/3T3 fibroblasts (ATCC® CRL-1658™) were maintained in DMEM (Gibco) supplemented with 10% FBS (Sigma Life Science), 1% Penicillin/Streptomycin (100 U/ml Penicillin, 100 µg/ml Streptomycin; Gibco), and 4.5 g/L α-D-glucose monohydrate (ROTH®). RMA and RMA-S, sublines of the B6 mouse-derived EL-4 T-cell leukemia, were maintained in RMPI (Gibco) supplemented with 10% FBS (Sigma Life Science), 1% Penicillin/Streptomycin (Gibco), 1% L-glutamine (Gibco), and 2-mercaptoethanol (Sigma).

Hoxb8-immortalized Cas9-progenitor cells were described in detail before (Hammerschmidt et al., 2018). Briefly, Hoxb8-immortalized Cas9-progenitor cells were cultured in complete IMDM (Merck) supplemented with 1% L-Glutamine, 1% penicillin/streptavidin, 10% heat-inactivated FBS, 100ng/ml mSCF, 100ng/ml huFLt3, 100ng/ml huIL-11, 20ng/ml mIL-3, 1µg/ml Doxycycline, and 1µg/ml Puromycin.

All cultures were passaged every second to the third day and kept at 37°C with 5% CO_2_.

### Hoxb8-immortalized Cas9-progenitor cell differentiation into macrophages

Hoxb8-immortalized Cas9-progenitor cells were differentiated into macrophages as described before (Hammerschmidt et al., 2018). Briefly, the cells were seeded in macrophage-differentiation medium consisting of IMDM (Gibco, with L-Glutamine) supplemented with 10% FBS (Sigma Life Science), 1% penicillin/streptomycin (Gibco), 150µM 1-thioglycerol (Sigma-Aldrich), and 5 ng/ml recombinant mouse macrophage colony stimulating factor (rm-M-CSF; PEPROTECH). The medium was exchanged on day 3 (retaining all cells) and day 6 (removing non-adherent cells). The cells were used for experiments on day 9 after initial seeding.

### In vitro *MCMV Infection of different cell lines*

For all infection experiments with RAW 264.7 and NIH/3T3 cell sublines, the cells were seeded at least 5 hours before the infection. If not otherwise indicated, we used a multiplicity of infection (MOI) 1 and incubated cells with viruses for 20-26 hr. Before infection, RMA and RMA-S cells were first maintained for 26-28 hrs at 26°C. Then, either 0.2% DMSO (v/v), 2 µg/ml MHC-I binding SIINFEKL peptide (ovalbumin 257-264, GenScript), or 2 µg/ml ISQAVHAAHAEINEAGR MHC-II binding peptide (ovalbumin 323-339, GenScript) was added. After additional 4 hours, cells were infected with MCMV-3D or -3DR at MOI 2 and maintained at 26°C for further 20-22 hours before analysis.

To detect the infected cells, we analyzed the percentage of mCherry+ cells on Cytek® Aurora spectral flow cytometer equipped with 355 nm, 405 nm, 488 nm, 561 nm, and 640 nm lasers. Flow cytometry data was then analyzed using FCS Express V7 (Denovo) or FlowJo V10 (BD). In some experiments, we also measured levels of Gaussia luciferase secreted into the supernatant by infected cells. For measurement, supernatants of cell culture media were diluted 1:9 with PBS and mixed with Gaussia luciferase substrate (final concentration of 0.1 μg Coelenterazine/ml). Bioluminescence was measured with SpectraMax® iD3 multi-mode microplate reader (Molecular Devices) using SoftMax Pro 7.03 SP1 software.

### Gene knockouts using CRISPR/Cas9 ribonucloparticles (RNPs)

For CRISPR/Cas9-mediated gene editing, RNPs consisting of Cas9 nuclease (Integrated DNA Technologies Inc.; IDT) and/or targeting guide RNA (gRNA; annealed crRNA and tracrRNA [IDT] or single-guide RNA [sgRNA; Synthego Inc.]) were used. The used crRNA and sgRNA sequences are listed in Supplementary Table 1. CrRNA:tracrRNA complexes were prepared and mixed with Cas9 as described (Durán et al., 2021). In experiments with sgRNAs, 210 pmol of each sgRNA was mixed with 70 pmol of Cas9, and incubated at room temperature for 10–20 min. Before use, all Cas9 RNPs were supplemented with Alt-R Cas9 Electroporation Enhancer (IDT).

For RNP delivery, we used SF Cell Line 4D-Nucleofector™ X Kit L (for Raw 264.7 cells) or SG Cell Line 4D-NucleofectorTM X Kit L (for NIH/3T3 cells; both from Lonza Inc.). Up to 5 × 10^6^ cells were resuspended in 100 μl of nucleofection solution, added to the cuvette pre-loaded with a total of 10 μl of RNPs, and immediately electroporated using pulse program DS136 (for Raw 264.7 cells) or EN158 (NIH/3T3 cells). Immediately after the nucleofection, cells were seeded in the pre-warmed medium.

The success of nucleofection was validated using flow cytometry. In cases when the protein of interest was still detectable, nucleofection was repeated up to four times per cell type and per RNP to reach a knockout score of >95%. In some cases, knockout cells were additionally sorted after labeling of remaining WT cells using appropriate antibodies.

### CRISPR/Cas9-mediated repair of B2m locus

We initially nucleofected Raw 246.7 cells with Cas9 RNPs targeting the *B2m* gene with crRNA 511-UGAGUAUACUUGAAUUUGAG-311 as described above. Afterward, cells were expanded for 2 days, and MHC-I-cells were FACS sorted. Next, cells were seeded for single-cell colonies that were, after expansion, screened using genome editing analysis. We selected only colonies carrying the type of mutation for repair with CRISPR/Cas9. Cells were then nucleofected with CRISPR/Cas9 RNPs containing 70 pmol of Cas9 complexed with 210 pmol of crRNA (5⍰-GCGUGAGUAUACUUGAAUUG-3⍰):tracrRNA together with 70 pmol of electroporation enhancer or with the same RNPs in the presence of 100 pmol of single-stranded DNA HDR template (5⍰-GTTTTCATCTGTCTTCCCCTGTGGCCCTCAGAAACCCCTCAAATTCAAGTATACTCACGCCACCCACCGGAGAATGGGAAGCC-3⍰). Electroporation was performed with prepared RNPs with or without HDR template with the same protocol for the knock-out experiment. After 7 days, we FACS sorted MCH-I+ cells using BD FACSAria™ Fusion cell sorter. Those cells, after expansion, were again seeded for single-cell colonies, which were then subjected to genome editing analysis. Only cells carrying integrated *B2m* gene template (corresponding to original gene sequence) were used in subsequent infection experiments.

### CRISPR/Cas9 genome editing analysis

To determine knockout or knock-in scores of CRISPR/Cas9-induced mutations, the DNA was first isolated using QIAamp DNA Mini Kit (Qiagen). The targeted gene region was amplified by PCR (NEB Next High-Fidelity 2xPCR Master Mix; New England Biolabs) using primers listed in Supplementary Table 2. Consecutively, the PCR products were purified using QIAquick® PCR Purification Kit (Qiagen) and sent for Sanger sequencing. Knockout or knock-in scores were determined from given sequences using the ICE analysis tool (https://ice.synthego.com/, Synthego).

### Cell line characterization and knockout analysis

We used flow cytometry to measure the protein expression of genes targeted with CRISPR/Cas9 genome editing. Unspecific antibody binding was blocked by a 15-minute incubation with mouse serum, rat serum, and/or homemade anti-CD16/CD32 antibodies. The staining using antibodies listed in Supplementary Table 3 was performed for 30 min at 4°C. After 2 washing steps, in some experiments, cells were additionally stained with DAPI (4,6-diamidino-2-phenylindole) for 2 minutes at room temperature. After extensive washing with PBS with 10% FCS, cells were acquired in an LSR II cytometer equipped with 405nm, 488 nm, and 633 nm lasers (BD). FCS files were analyzed using FCS Express V7 (De Novo Software) or FlowJo V10 (BD).

### Genome-wide CRISPR-Cas9 knockout screen

To screen for genes enabling MCK2-mediated MCMV infection, we first generated SpCas9 and the blasticidin resistance gene expressing *Nrp1*^*-/-*^ NIH/3T3 fibroblasts. We seeded 5 × 10^4^ cells per well of a 12-well plate and transduced them the next day with 0.005 µL of 100-fold concentrated lentiviral LKO5.SF.SpCas9.P2A.BSD.PRE supernatant. Transduction was supported by 4 µg/ml protamine sulfate and centrifugation for 1 hour at 863g and 37°C. Subsequently, transduced cells were selected with 10 µg/mL blasticidin (invivoGen) for 13 days. Next, SpCas9-expressing *Nrp1*^*-/-*^ NIH/3T3 fibroblasts were then transduced with the Mouse Brie CRISPR knockout lentiviral prep (Addgene #73633-LV) as described previously (Doench et al., 2016; Sanson et al., 2018). Next, two biological replicates of cells expressing mouse Brie library were infected with MCK2^+^ MCMV-3DR at MOI 10. Infected samples were purified for mCherry^-^ cells at 20h.p.i. by FACS, and re-infected at MOI 10 on the same day. The next day the mCherry^-^ cells were also sorted, and expanded for 2 days before the 3rd infection with MCMV-3DR at MOI 16. Again, at 20 h p.i. mCherry^-^ cells were FACS sorted. For one biological replicate we used those cells for DNA isolation, while the second biological replicate was expanded for another 2 days, infected a 4th time with MCMV-3DR at MOI 16, and mCherry^-^ cells at 20 h p.i. were used for DNA extraction.

For both biological replicates as well as from the uninfected input cells, the genomic DNA was extracted, sgRNAs were amplified by PCR, and purified using the AMPure XP-PCR purification system according to the protocol provided by BROAD Institute (https://media.addgene.org/cms/filer_public/61/16/611619f4-0926-4a07-b5c7-e286a8ecf7f5/broadgpp-sequencing-protocol.pdf). Prepared libraries were sequenced using NextSeq 550 (Illumina) and generated FASTQ files were analyzed using a Bioconductor package edgeR in R (Robinson et al., 2010). Selected genes were chosen if they had a count per million (CPM) count >5 in all 3 samples analyzed (sample going through MCMV-selection vs. 2 biological replicates of input cells). The analysis was done according to the edgeR user guide for CRISPR-Cas9 screen analysis (version May 12th, 2021; https://bioconductor.org/packages/release/bioc/vignettes/edgeR/inst/doc/edgeRUsersGuide.pdf).

### Experimental animals

C57BL/6N, BALB/c, and C3H mice were purchased from Charles River, while B6.129P2-*B2m*^*tm1Unc*^/J mice, homozygous for the *B2m*^*tm1Unc*^ targeted mutation, and B6.129P2(Cg)-Cx3cr1^tm1Litt^/J, CX3CR1^GFP^ knock-in/knock-out mice were from The Jackson Laboratory. All mice were bred and maintained in the Central Animal Facility of Hannover Medical School (Hannover, Germany) under specific pathogen-free conditions and used for experiments at the age of 7 – 26 weeks. All experiments involving mice were handled in compliance with the European and national regulations for animal experimentation (European Directive 2010/63/EU; Animal Welfare Acts in Germany) and were approved by the Niedersächsisches Landesamt für Verbraucherschutz und Lebensmittelsicherheit (LAVES; Lower Saxony, Germany).

### Ex vivo *infection of alveolar macrophages*

Cells were collected by broncho-alveolar lavage (BAL) through a plastic catheter clamped into the trachea using three separate lavage steps with 400 µl, 300 µl, and 300 µl of PBS each. On average >95% of cells in BAL samples from uninfected mice are alveolar macrophages (Bošnjak et al., 2019, 2021). Afterwards, the cells were counted using Neubauer chambers, centrifuged for 10 min at 300 g at 4°C, plated on 96-well plates in RPMI media, and subsequently infected with indicated MOI of MCMV mutants. At 20-24 h p.i., cells were collected using 0.25% trypsin (Biochrom AG, Berlin, Germany) for the detachment and mCherry expression indicating infection with the viruses was analyzed using flow cytometry. Alveolar macrophages were identified by forward and side scatter position and autofluorescence.

### Intranasal infection with MCMV

For infection with the indicated MCMV strain, we intranasally (i.n.) administered 10^6^ plaque forming units (PFU) of MCMV resuspended in 40 µl PBS to the nostrils of lightly anaesthetized mice.

### Sample collection from mice

From uninfected mice and mice at 1or 7 days post-infection blood was collected from inferior vena cava into K3 EDTA tubes (Sarstedt). Afterward, salivary glands, lungs, left kidney, median liver lobes, and spleens were resected. One salivary gland was used for virus titer determination and luciferase assay, while the second salivary gland was longitudinally divided into two halves: one for DNA extraction and the second for flow cytometry analysis.

From the lungs, the post-caval lobe was used for DNA extraction, the middle lobe for virus titer determination and luciferase assay, the superior and inferior lobes for flow cytometry, and the left lung lobe for immunohistology.

### Virus titers and luciferase assay

The viral plaque formation and luciferase assay were performed as described earlier (Lueder et al., 2018). Briefly, explanted organs were stored in 600 μL DMEM containing 10% FCS and 1% Penicillin/Streptomycin (Gibco). After mechanical disruption by shaking at 25/s over 4 min with metal beads (TissueLyser II, Qiagen), two 250 μL aliquots of the homogenate were stored at -80°C for viral plaque titration. The remaining homogenates were centrifuged at maximum speed for 15 min and used to prepare 180 μL of a homogenate 1:10 dilution in PBS. Then, 20 μL of the substrate (1 μg Coelenterazine/ml in PBS) was added and luminescence was measured for 10s in Lumat LB 9507 (Berthold Technologies) in duplicates. The viral plaque titration was performed on MEF monolayers in duplicates (Lueder et al., 2018).

### Immunohistology

The left lung lobe was filled via the bronchus with PBS solution containing 2% paraformaldehyde and 30% sucrose. Lungs were incubated in 500 µl of the same solution overnight. Fixed lungs were embedded in OCT compound (TissueTek, Sakura), frozen at -20°C, and cut in 8 µm thick cryosections, which were blocked with rat serum in TBST and stained with anti-CD11c-PE-Cy7 (Supplementary Table 3) and DAPI. Images were taken with AxioScan.Z1 (Carl Zeiss) in combination with the Colibri 7 (Carl Zeiss) as well as Plan-Apochromat objectives 20x/0.8 (Magnification/numerical aperture) and processed with ZEN blue 3.2 software (Carl Zeiss). Only sections including the main bronchus were used for analysis, and for each animal one lung section was analyzed. Infected cells were identified by mCherry expression, morphology, and DAPI staining and further divided into alveolar macrophages (identified by CD11c expression and localization in the alveolar space) or other cells.

### Immunophenotyping of infected cells and leukocyte infiltrates in the lungs or salivary glands using spectral flow cytometry

For flow cytometry, right lung lobes were inflated with digestion medium # 1 made of RPMI without phenol red (Gibco) supplemented with 5% of FCS, 4 U/ml of Elastase (Worthington Biochemical Corporation), 1U/ml of Dispase II (Sigma), and 200 µg/ml of DNase I (Sigma). After incubation in wells containing 2 ml of digestion medium # 1 for 45 minutes at 37°C, lung lobes were minced into small pieces using sharp scissors. Tissue pieces then were transferred to a 50-mL conical tube containing 5 ml of digestion medium #2 (RPMI without phenol red containing 5% of FCS, 25 µg/ml of Liberase (Roche), and 200 µg/ml of DNase I (Sigma)). After the second incubation for 30 minutes at 37 °C, single lung cells were then separated by smashing samples through 40 μm cell strainers. The remaining RBCs were removed using an erythrocyte lysis buffer and single-cell suspensions were prepared by filtering again through 40 μm cell strainers.

Isolated cells were blocked with 10% rat serum for 10 minutes at 4°C before antibody mixes were added, and cells were incubated for additional 30 minutes at 37°C. The antibodies used for staining are listed in panel 1 in Supplementary Table 3. After washing twice, samples were acquired using the Cytek Aura flow cytometer (Cytek) which is equipped with 355 nm, 405 nm, 488 nm, 561 nm, and 640 nm lasers. Flow cytometry data was then analyzed using FCS Express V7 (Denovo) or FlowJo V10 (BD).

### Statistical Analysis

Statistical analysis was performed with GraphPad Prism 8 using Welch’s ANOVA and Tukey’s multiple comparisons test for comparison of groups with equal standard deviations and Dunnett’s T3 multiple comparisons test for experiments with variation among the standard deviations. Where indicated, we logarithmically transformed the data before analysis. P-values <0.05 were considered significant.

