## Supplemental Figures 1-10 and Tables 1-3 for "MHC class Ia molecules facilitate MCK2-dependent MCMV infection of macrophages and virus dissemination to the salivary gland"

* This authors equally contributed

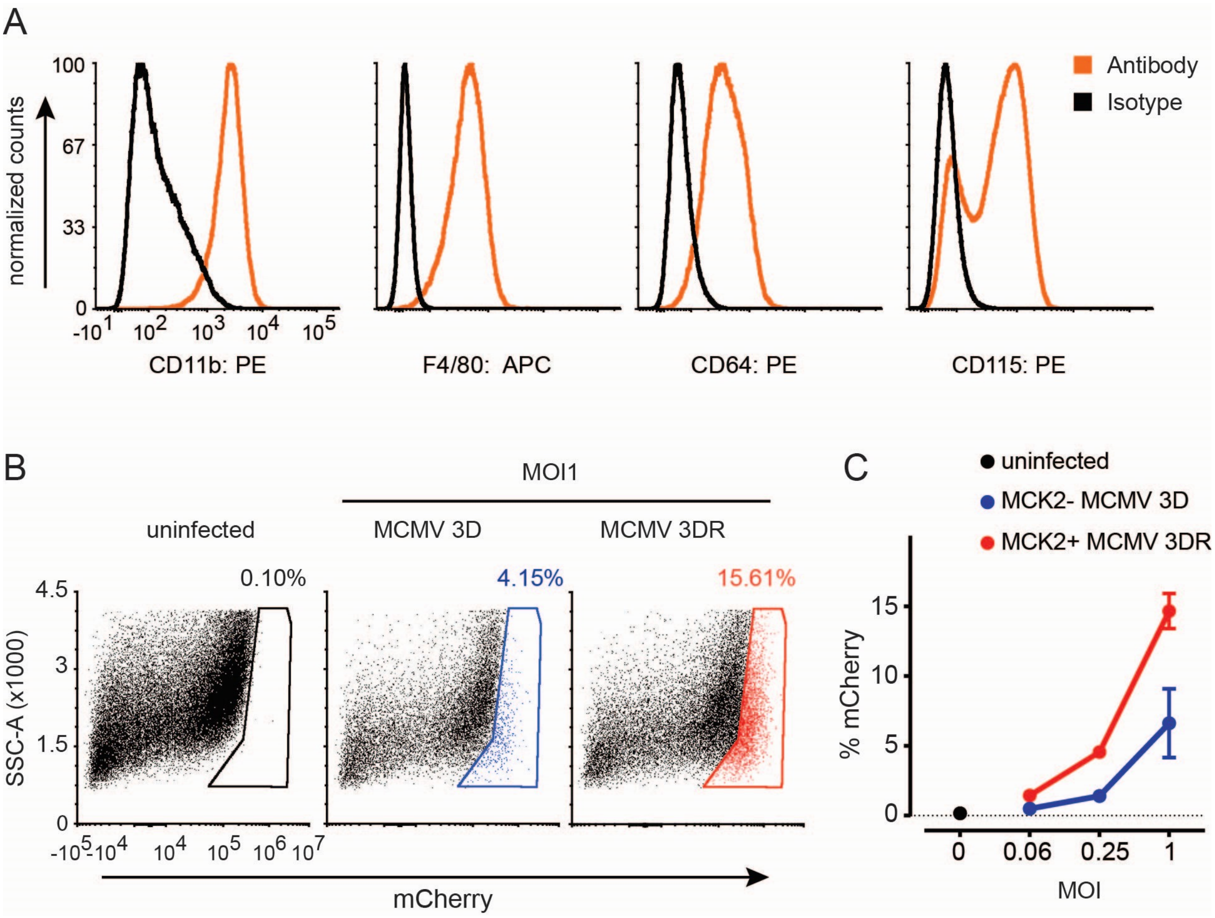


**Supplementary Figure S1 (related to Figure 1). MCMV infection of Cas9-Hoxb8 derived macrophages depends on MCK2.**

**(A)** Phenotype of Cas9-Hoxb8 derived macrophages after 9 days of differentiation in medium containing recombinant murine macrophage colony stimulating factor (rm-M-CSF).

**(B)** Representative flow cytometry dot plots of uninfected Cas9-Hoxb8 derived macrophages and cells at 18-22 hours post infection with indicated MCMV strains at MOI of 1.

**(C)** Quantification of infected cells measured as percentage of mCherry positive Cas9-Hoxb8 derived macrophages by flow cytometry at 20 h p.i. with indicated doses of MCMV-3D or MCMV-3DR. Representative data from two independent experiments. Dots represent the mean of a technical triplet, error bars are SEM.

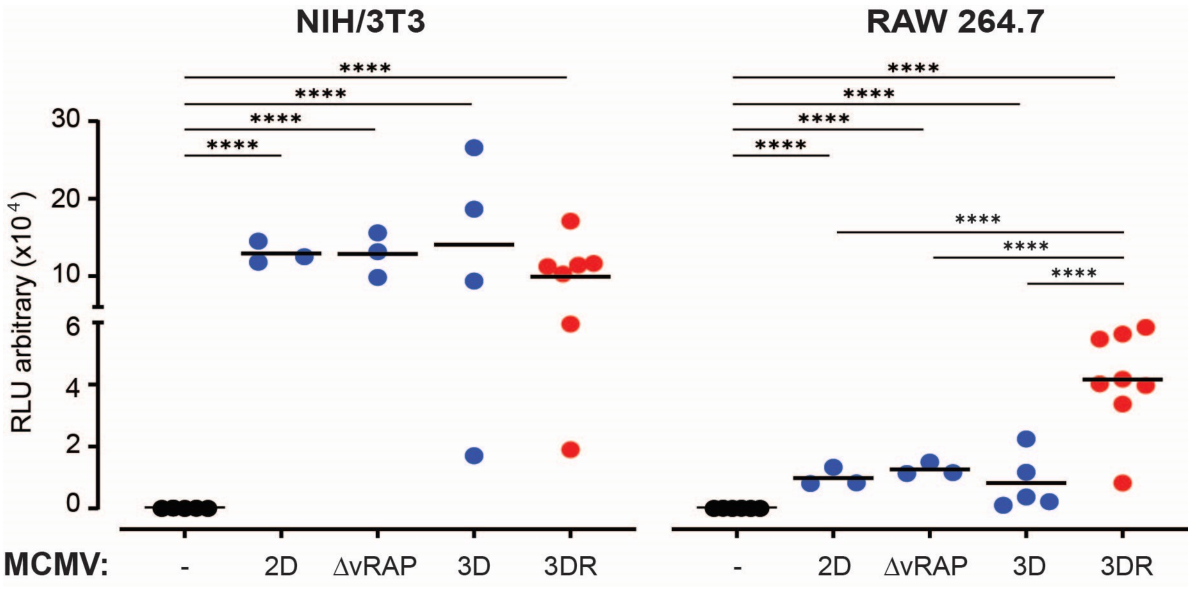


**Supplementary Figure S2 (related to Figure 1).** Gaussia luciferase activity in supernatant of NIH/3T3 fibroblasts and RAW 264.7 monocyte/macrophage cell cultures at 18-22 hr. p.i. with indicated MCMV strains at multiplicity of infection (MOI) of 1. One dot represents the mean of a technical triplet, data are pooled from 3-7 independent experiments, and the group means are shown as lines. Statistical Analysis: Welch's ANOVA test followed by Dunnett's T3 multiple comparison test; *p < 0.05; ****p < 0.0001.

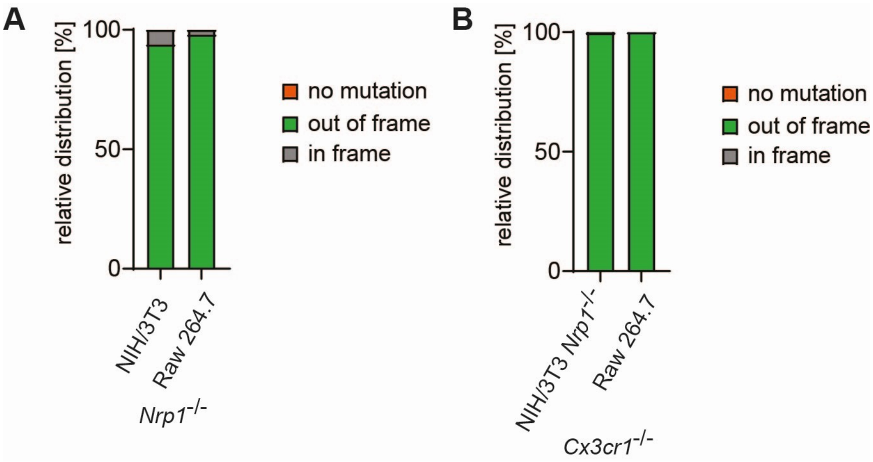


**Supplementary Figure S3 (related to Figure 2).** Relative distribution of mutations in cells nucleofected with CRISPR/Cas9 ribonucleoparticles (RNPs) targeting **(A)** *Nrp1* or **(B)** *Cx3cr1*.

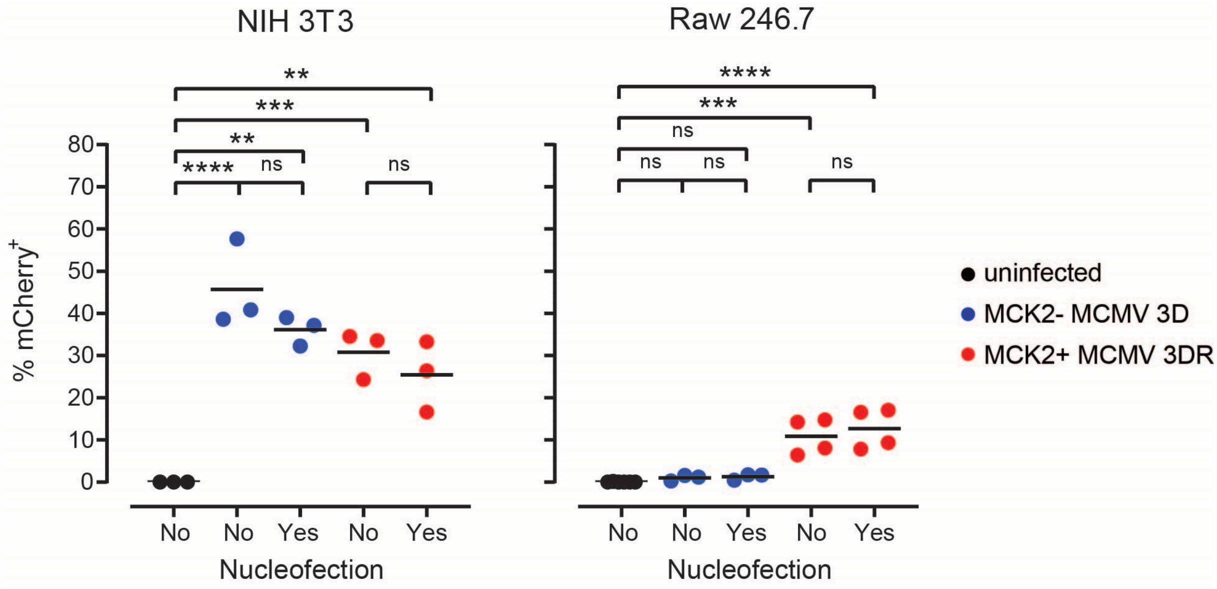


**Supplementary Figure S4 (related to Figure 2).** Quantification of mCherry signal at 18-22 h p.i. with indicated MCMV strains at multiplicity of infection (MOI) of 1 from untreated cells (No nucleofection) and cells treated with control CRISPR/Cas9 ribonucleoparticles (Yes nucleofection). Data are from 3-4 independent experiments. One dot equals a mean of the technical triplicates from one experiment, line at mean value per group. Statistical analysis: One-way ANOVA test followed by Sidak's multiple comparison test; ns, not significant; ** $p < 0.01$, *** $p < 0.001$, ****$p < 0.0001$. Data from nucleofected and infected cells are identical to that shown in Fig. 2c.

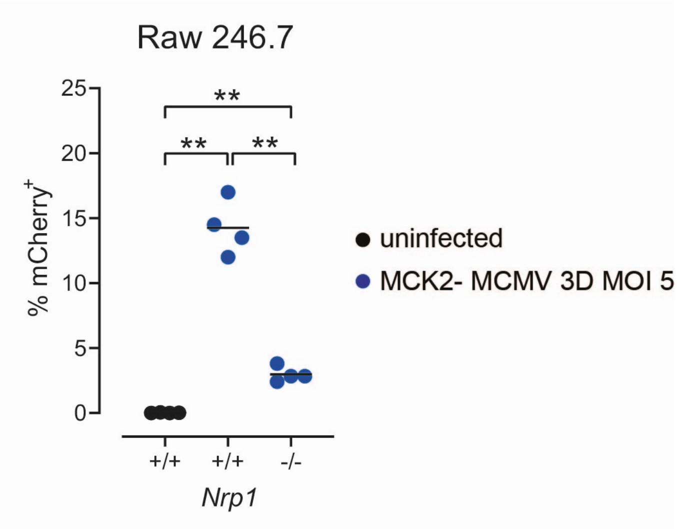


**Supplementary Figure S5 (related to Figure 2).** Quantification of mCherry signal at 18-22 h p.i. with MCMV-3D at multiplicity of infection (MOI) of 5 from cells treated with Ctrl. RNPs or Nrp1 RNPs. Data are from 4 independent experiments. One dot equals a mean of the triplicates from one experiment, line at mean value per group. Statistical analysis: Welch's ANOVA test followed by Dunnett's multiple comparison test; ** p < 0.01.

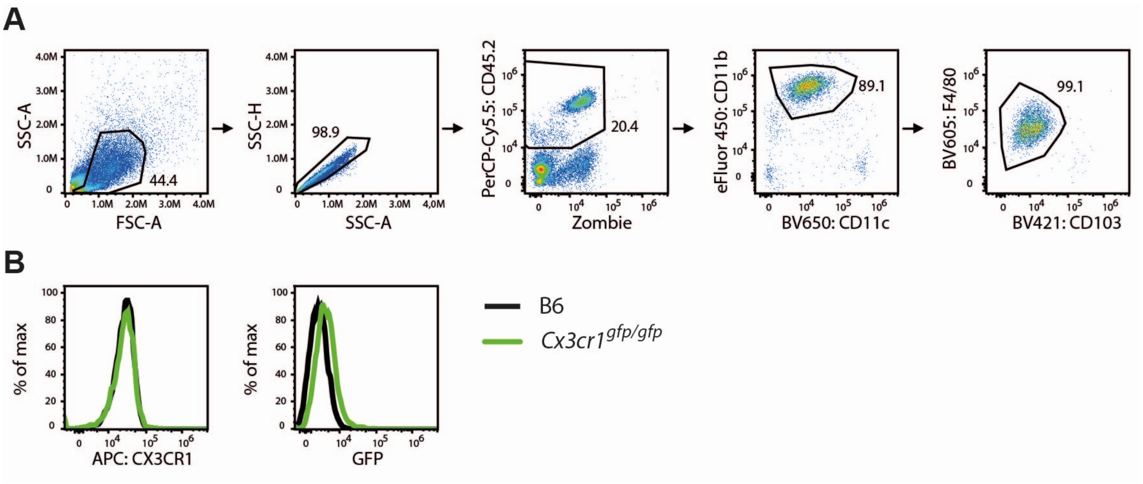


**Supplementary Figure S6 (related to Figure 2).**

(A) Gating strategy for detection of alveolar macrophages within broncho-alveolar lavage stained with antibody panel 2 listed in Supplementary Table 3.

(B) Representative histograms from wild-type B6 and CX3CR1$^{gfp/gfp}$ mice indicating staining for CX3CR1 and GFP that was knocked-in instead of *Cx3cr1*.

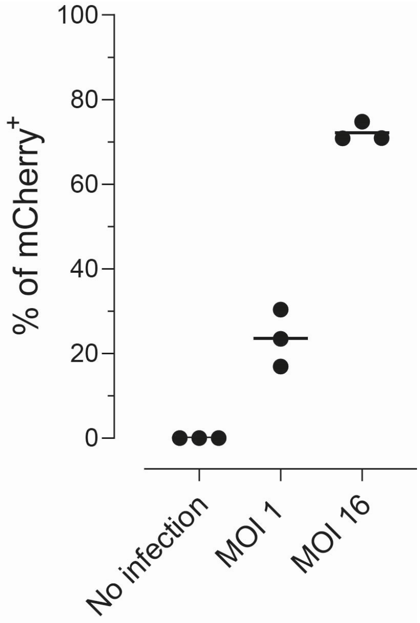


**Supplementary Figure S7 (related to Figure 3).**

Susceptibility of *Nrp1*$^{-/-}$ NIH/3T3 fibroblasts towards MCMV-3DR at multiplicities of infection (MOI) as indicated. Cells were analysed at 22 h p.i. Data are from one experiment, each dot represents a value from an independent well.

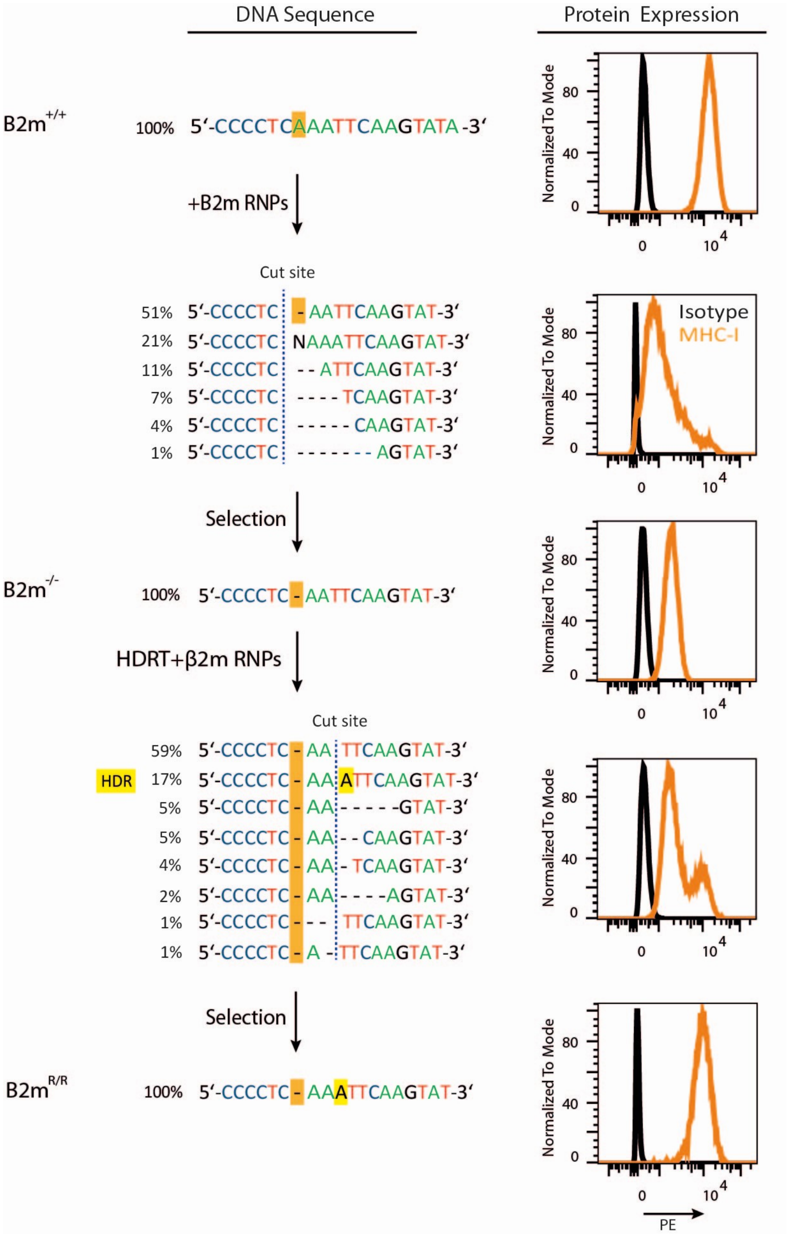


**Supplementary Figure S8 (related to Figure 4).**

MHC-I expression in Raw 246.7 cells before manipulation, after treatment with beta-2 microglobulin (B2m) ribonucleoparticles (RNPs), selection of *B2m*$^{-/-}$ cells by single cell colonies, expansion, repair of *B2m* locus via CRISPR/Cas9 gene editing using homology-directed repair (HDR) template (HDRT), and selection of cells with correctly repaired *B2m* locus (*B2m*$^{R/R}$) by single cell colonies. For details, please refer to Materials and methods.

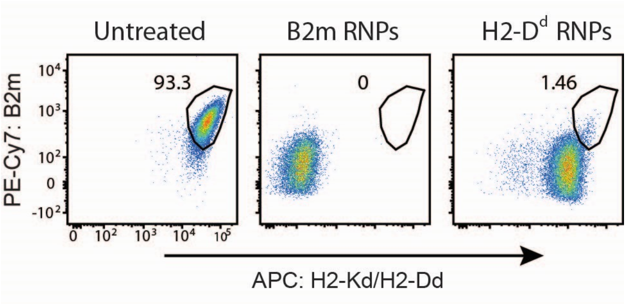


**Supplementary Figure S9 (related to Figure 5).**

Representative flow cytometry dot plots of showing expression of H2-Kd/H2-Dd (antibody clone 34-1-2S) and beta-2 microglobulin (B2m) on untreated Raw 264.7 cells and cells nucleofected with ribonucleoparticles (RNPs) targeting *b2m*, *H2-K1*, or *H2-D1*. Data are from representative of two independent experiments.

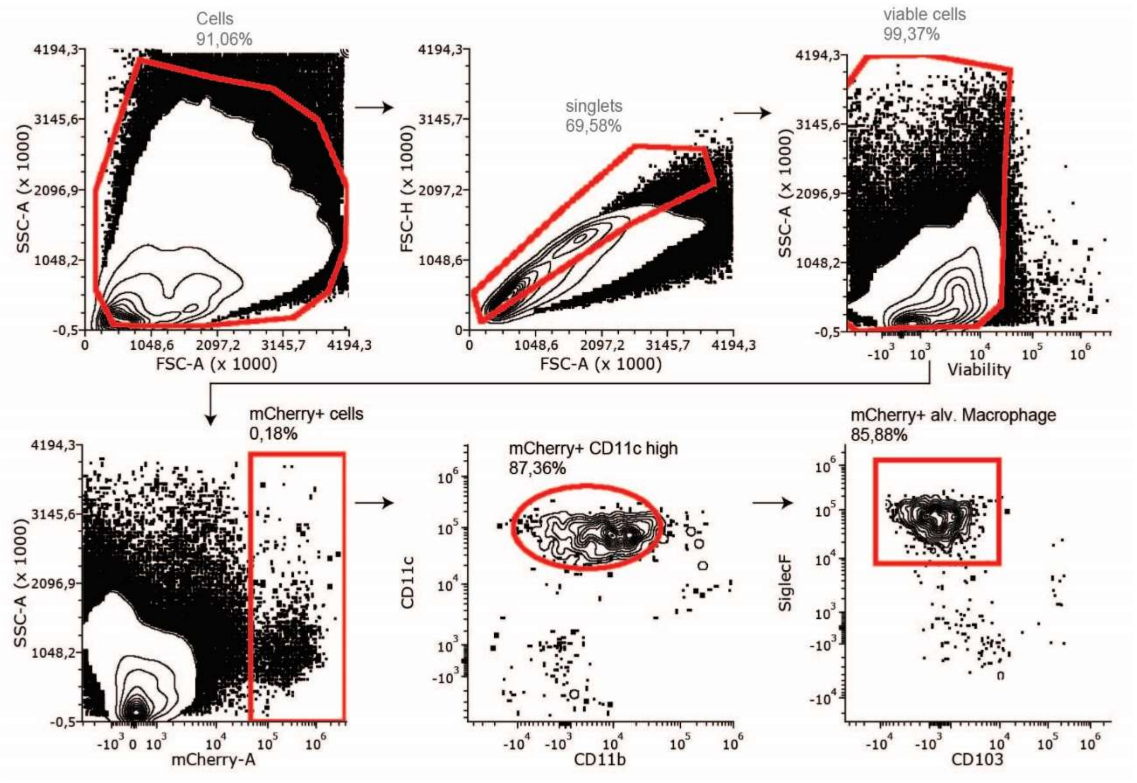


**Supplementary Figure S10. Gating strategy of alveolar macrophages in the lungs of mice infected with MCMV-3DR or -3D (related to Figure 6).**

Density plots showing gating strategy for mCherry$^{+}$ alveolar macrophages in the lungs analyzed with antibody panel 1 listed in Supplementary Table 3. Data shown are from a representative mouse at 1 day post intranasal infection with MCMV-3DR.

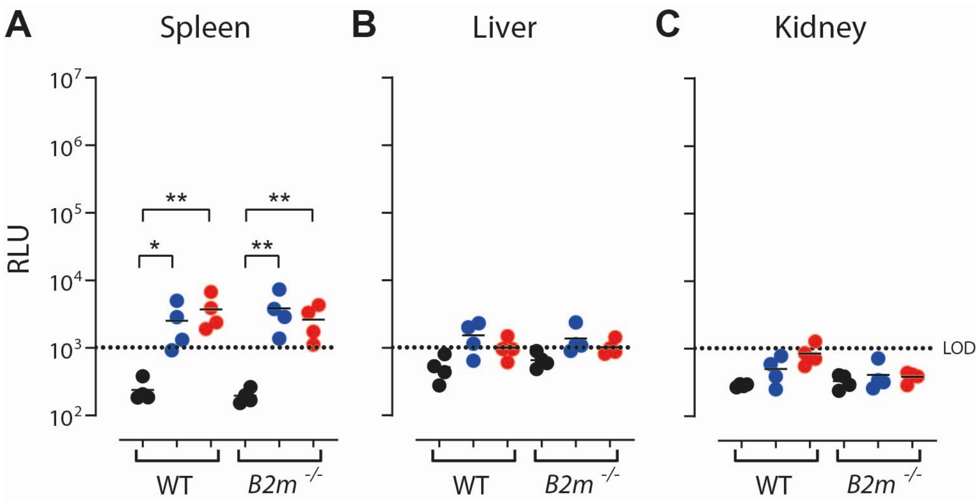


**Supplementary Figure S11 (related to Figure 7).**

Luciferase activity in **(A)** spleens, **(B)** livers, and **(C)** kidneys of wild-type B6 (WT) or *B2m*$^{-/-}$ mice 7 days post intranasal infection (dpi) with MCMV-3D or -3DR. LOD – limit of detection.**(A-C)** Pooled data from two experiments with 4 mice per group. Statistical analysis: Welch's ANOVA test followed by Dunnett's T3 multiple comparison test on log-transformed values; * $p < 0.05$; ** $p < 0.01$.

**Table S1: Related to STAR methods. List of crRNAs.**

| *Gene target* | *crRNA sequence (5‘->3‘)* | *Sequence origin* | *Comment* |
|---|---|---|---|
| *nrp1* | CACAAUAACGCCCAAUGUGA | IDT | Knock-out |
| *b2m* | CCUCACAUUGAAAUCCAAAU | Brie library, Addgene | Knock-out |
| | GUAUGCUAUCCAGAGUGAGU | Brie library, Addgene | Knock-out |
| | CUCAAAUUCAAGUAUACUCA | Brie library, Addgene | Knock-out |
| | UGAGUAUACUUGAAUUUGAG | IDT | Knock-out for repair |
| | GCGUGAGUAUACUUGAAUUG | IDT | Knock-in |
| *cd81* | AGGUGUAGCUCUGUGGUUGC | Brie library, Addgene | Knock-out |
| | GGGCUUCGUAAACAAAGACC | Brie library, Addgene | Knock-out |
| | UUCAGCAAGCUGUGAUGGAU | Brie library, Addgene | Knock-out |
| *cx3cr1* | CAGACCGAACGUGAAGACGA | Syntego Inc. | Knock-out |
| | UUGGGCGACAUUGUGGCCUU | Syntego Inc. | Knock-out |
| | AAAUUCUCUAGAUCCAGUUC | Syntego Inc. | Knock-out |
| *H2-K1* | CGCUGAUCACCAAACACAAG | IDT | Knock-out |
| | GGUGACUUCACCUUUAGAUC | IDT | Knock-out |
| *H2-D1* | UGUCGGCUAUGUGGACAACA | IDT | Knock-out |
| | GGUGACUUCACCUUUAGAUC | IDT | Knock-out |
| | GCAUUACAAGGCCUACCUGG | IDT | Knock-out |

**Table S2: Related to STAR methods. List of primers used**

| **Target gene** | **primer type** | **primer sequence (5' -> 3')** |
|---|---|---|
| *nrp1* | forward primer | ATTGTCCTGCAAATGCTCCTC |
| *nrp1* | reverse primer | CCTGACACCTGTGCCTTTGA |
| *cX3cr1* | forward primer 1 | GAAGAAGGCAGTCGTGAGCT |
| *cx3cr1* | reverse primer 1 | CACCCTTTCAGTGTTTTCTCCC |
| *cx3cr1* | forward primer 2 | CAGTCGTGAGCTTGCACATG |
| *cx3cr1* | reverse primer 2 | CACCCTTTCAGTGTTTTCTCCC |
| *b2m* | forward primer | GACACTGCTAAAAGCCAGGT |
| *b2m* | reverse primer | CAGATGGAGCGTCCAGAAAGT |

**Table S3: Related to STAR methods. List of antibodies and other reagents used for cell line characterization and knockout analysis.** Panel 1 – Staining of lung cells, Panel 2 – staining of broncho-alveolar lavage cells; NA – not applicable

| Antibody | Clone | Source | Identifier | Panel |
|---|---|---|---|---|
| Analysis of cell cultures | | | | |
| PE/Cy7 anti-mouse CD304/ Neuropilin-1 | 3E12 | BioLegend | Cat# 145212; RRID: AB_2562360 | NA |
| APC anti-mouse CX3CR1 | SA011F11 | BioLegend | Cat# 149008; RRID: AB_2564492 | NA |
| APC anti-mouse CX3CR1 | QA16A03 | BioLegend | Cat# 153708; RRID:AB_2734224 | NA |
| PE anti-mouse H-2Kd | SF1-1.1 | Thermo Fisher Scientific | Cat# 12-5957-82; RRID:AB_2043875 | NA |
| FITC anti mouse H-2Kk | 36-7-5 | BioLegend | Cat# 114905; RRID:AB_313612 | NA |
| FITC anti mouse H-2Kq | KH114 | BioLegend | Cat# 115104; RRID:AB_313625 | NA |
| PE anti-mouse H-2Kk antibody | 36-7-5 | BioLegend | Cat# 114907; RRID:AB_313614 | NA |
| APC anti-mouse H-2Kd/H-2Dd antibody | 34-1-2S | BioLegend | Cat# 114714; RRID:AB_2734174 | NA |
| PE/Cyanine7 anti-mouse beta2-microglobulin antibody | A16041A | BioLegend | Cat# 154508; RRID:AB_2728213 | NA |
| Analysis of cells isolated from lungs and broncho-alveolar lavage | | | | |
| BV421 anti-mouse CD103 antibody | 2E7 | Biolegend | Cat# 121422, RRID:AB_2562901 | 1, 2 |
| eF450 anti-mouse CD11b antibody | M1/70 | Thermo Fisher Scientific | Cat# 48-0112-82, RRID:AB_1582236 | 1, 2 |
| BV605 anti-mouse F4/80 antibody | BM8 | Biolegend | Cat# 123133, RRID:AB_2562305 | 2 |
| BV650 anti-mouse CD11c antibody | N418 | Biolegend | Cat# 117339, RRID:AB_2562414 | 1, 2 |
| PerCP-Cyanine5.5 anti-mouse CD45.2 antibody | 104 | Thermo Fisher Scientific | Cat# 45-0454-82, RRID:AB_953590 | 2 |
| APC-Cy™7 Rat Anti-Mouse Siglec-F antibody | E50-2440 | BD Biosciences | Cat# 565527, RRID:AB_2732831 | 1 |

| | | | | |
|---|---|---|---|---|
| APC anti-mouse CX3CR1 | SA011F11 | BioLegend | Cat# 149008; RRID: AB_2564492 | 2 |
| Zombie NIR™ Fixable Viability | N/A | Biolegend | N/A | 1, 2 |
| Antibodies used for histology | | | | |
| PE-Cy7 anti-mouse CD11c antibody | N418 | Biolegend | Cat# 117318, RRID: AB_493568 | NA |
